# Temperature variation and life history mediate the degree of nonlinearity in fluctuations of global marine fish populations

**DOI:** 10.1101/2025.07.13.664566

**Authors:** Robert M. Hechler, Martin Krkosek

**Affiliations:** Department of Ecology and Evolutionary Biology, University of Toronto; Toronto, M5S3B2, Canada

**Keywords:** Fisheries, nonlinear dynamics, physical-biological interactions, empirical dynamic modeling, noise amplification

## Abstract

Nonlinear dynamics readily occur in natural ecosystems and can drive irregular population fluctuations through oscillations, chaos and alternative stable states. However, the effects of anthropogenic changes, such as to demography and the climate, on nonlinearity of population fluctuations is unknown. We evaluated the extent and magnitude of nonlinearity and its environmental and life history correlates in 243 recruitment and 267 spawner time series of 143 marine fish species, worldwide. Here we show that temperature variation amplifies nonlinearity in recruitment and spawner biomass, while life history mediates the degree of nonlinearity for the latter, dampening it in slow-lived species. Nonlinearity was displayed by 81% of populations and correlated with the magnitude of fluctuations. These nonlinear dynamics were low-dimensional and causally forced by temperature in 60% of populations with the probability of forcing increasing for recruits in variable temperature environments and fast-lived spawners. Our results challenge assumptions of stable dynamics and sustainable yield common to fisheries management, and suggests nonlinear fluctuations of fish populations are magnified by size-selective fisheries and environmental variability from global climate change.

## Main

Animal populations often show irregular fluctuations in abundance or biomass that are driven by deterministic nonlinear population dynamics such as sustained oscillations, alternative stable states and chaos^1–5^. Such complex behaviours are widespread in nature^3,6^ and arise from nonlinearity, where outputs do not change proportionally with inputs as relationships between governing variables vary with the system’s state. The degree of nonlinearity is correlated with the coefficient of variation in fluctuations^7,8^ and is influenced by intrinsic and extrinsic factors^6,8,9^. Importantly, intrinsic nonlinear fluctuations may be amplified by extrinsic environmental forcing, which is distinct from random fluctuations around an equilibrium that track environmental stochasticity. Understanding causes of nonlinearity is crucial for predicting and managing populations in the Anthropocene due to human-induced changes to demography^10^ and the climate^11^. However, the relative contributions of intrinsic and extrinsic factors to nonlinearity of population dynamics remains contentious and unclear.

Early theory demonstrated that nonlinearity can arise from intrinsic mechanisms such as increasing growth rates and delayed density dependence^1^. In the absence of external disturbances, nonlinear dynamics like chaos readily occur in experimental plankton communities^4^, flour beetle populations^12^, and single-species protist systems^13^. Chaotic fluctuations were found in >30% of 172 time series predominately from birds and mammals^3^. Further analysis of 502 insect, 53 fish, 51 bird, and 36 mammal time series showed that the likelihood of displaying nonlinear dynamics increased for species with fast life-history traits^6^. However, marine fish were only sparsely represented^6^ and empirical evidence linking life history traits and the magnitude of nonlinearity is lacking.

Marine fishes have long been studied for the roles of harvesting and environmental variability in driving fluctuations. Increased variability in abundance of harvested populations was first shown in a single-species logistic growth model^14^ and empirically confirmed using time series of exploited and unexploited larval fish^15^. Two competing hypotheses have been presented which are associated with fishing-induced age-truncation as harvesting removes older and larger individuals^10^. The external forcing hypothesis predicts a linear signature in the time series of exploited populations as smaller, younger fish are less able to smooth out environmental effects and track fluctuations more directly^8,10,14^. In contrast, the noise amplification hypothesis predicts an enhanced nonlinear signature in exploited populations due to interactions between nonlinearities and process noise^8,16^. There is growing evidence supporting the latter as multiple fish populations have been found to display nonlinear dynamics^8,9,16–21^ which can be harnessed to produce more accurate forecasts than conventional stock assessment models^21^ and even the true data generating model^22^.

Environmental variability can provide noise that interacts with nonlinearities to sustain population oscillations that would otherwise decay^23^. For example, episodic fluctuations in a reef fish result from amplification of wind stress by nonlinear processes in the larval phase^16^. Furthermore, stability analysis of a parametric model fitted to marine fish populations suggested that most populations were in the stable region of parameter space, but with damped oscillations that produced nonlinear dynamics due to the interaction with process noise^2,24^. The same model fitted to time series of *Daphnia* from a fully crossed experiment of environmental stochasticity and size-selective harvest indicated that nonlinearity resulted from the interaction of damped oscillations with demographic noise rather than process noise^25^. Time series of larval fish in California indicate that exploited marine fishes exhibit amplified nonlinearity which is hypothesized to result from fishing-induced age truncation that alters demographic parameters and interactions with process noise^8,15^. However, evidence regarding the extent of nonlinear noise amplification in natural populations is lacking and the acting mechanism remains unclear. More broadly, the role of the environment in driving the degree of nonlinearity and the extent to which nonlinearity varies with life history traits has not been explored.

Here, we investigate the extent of nonlinear dynamics in global marine fish populations, and the environmental and life history correlates of the degree of nonlinearity. We extracted 267 spawner biomass and 243 recruitment time series across 143 species from the RAM Legacy Stock Assessment Database^26^ (Fig. 1A and fig. S1). Additionally, we obtained correlates such as species life history traits^27^ (fig. S2) and sea surface temperature (SST) time series^28^ using delineated population boundaries^29^. We used methods from empirical dynamic modeling to analyze the time series^30^. We used simplex projection^31^ to determine the embedding dimension (E), which is the number of time lags needed to reconstruct the system in the state-space and provides an estimate of complexity by indicating how many processes are interacting to produce the observed dynamics. We then quantified the degree of nonlinearity for each population using sequentially weighted global linear map (S-map) and selected the nonlinear tuning parameter, θ, that maximized the out-of-sample prediction skill, ρ, using leave-one-out cross-validation. At θ = 0, the S-map is a global linear model that weights all points on the attractor equally, but as θ > 0 the model becomes nonlinear and more weight is given to points nearby on the attractor^32^. We used Bayesian models to estimate associations between several SST and life history correlates with dimensionality (E), prediction skill (ρ), and two metrics of nonlinearity that have been previously used in comparative analyses (𝜃_ρ_*max*__ and 𝛥𝜌 = 𝜌_*max*_ − 𝜌_θ=0_)^2,7–9,16^. We further investigated the importance of SST as a causal driver of fish dynamics using convergent cross mapping^33^ and then applied Bayesian models to estimate how the probability of temperature forcing on nonlinearity varies with SST, life history and habitats.

**Fig. 1.**
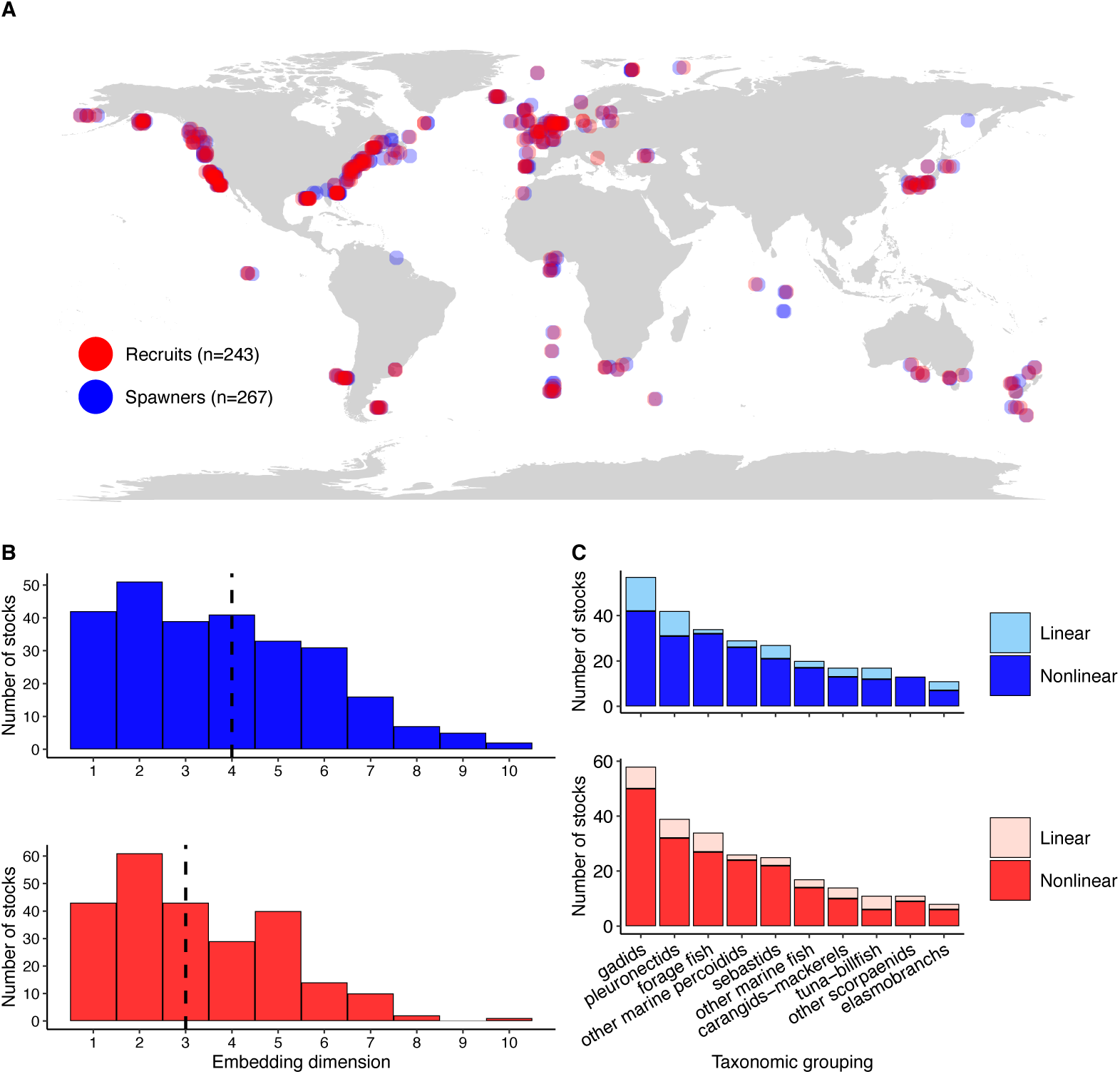
Marine fish populations are governed by low-dimensional, nonlinear dynamics. **(A)** Points represent the mean latitude and longitude of population boundaries for time series of recruits (abundance) and spawners (biomass) used in our analysis. **(B)** Histogram of the optimal embedding dimension (E) of fish populations with dashed lines showing the median E values. **(B)** Nonlinear dynamics are displayed by most recruit (201/243) and spawner (214/267) populations as identified by S-map nonlinear tuning parameter θ > 0 and Δρ >0.

## Results

We found that most marine fish populations displayed nonlinear dynamics (Fig. 1C). The degree of nonlinearity was mediated by SST variation and life history, depending on life stage (Fig. 2, figs. S3-7 and Table 1). For recruitment, nonlinearity became greater with increasing intra-annual SST variation (Fig. 2B; 95% credible interval (CI) for θ = 0.13 to 0.73 and Δρ = 0.07 to 0.52) and there was no clear effect of the other SST predictors or life history traits, as indicated by posterior distributions centered near 0 and 95% CIs that largely overlap 0 (Fig. 2). Nonlinearity of spawner biomass increased with the first principal component of an ordination of life history traits (PC1), representing fast life history traits like fast somatic growth and young age at maturity (Fig. 2D; 95% CI for θ = 0.07 to 0.52 and Δρ = 0.26 to 0.74) and negative values of principal component 2 (PC2), representing high fecundity (95% CI for θ = -0.55 to -0.01 and Δρ = -0.70 to -0.12). Nonlinearity of spawner biomass was also affected by mean SST (95% CI for θ = -0.46 to 0.03 and Δρ = -0.45 to 0.10), inter-annual SST variation (95% CI for θ = -0.30 to 0.21and Δρ = -0.53 to 0.05) and intra-annual SST variation (95% CI for θ = 0.13 to 0.63 and Δρ = 0.17 to 0.71) (Figs. 2A and 2C). These results were consistent across two metrics of nonlinearity (Fig. 2), and five sensitivity analyses that considered two SST datasets, accounted for uncertainty in θ and corrected for phylogenetic non-independence (figs. S3-7).

**Fig. 2.**
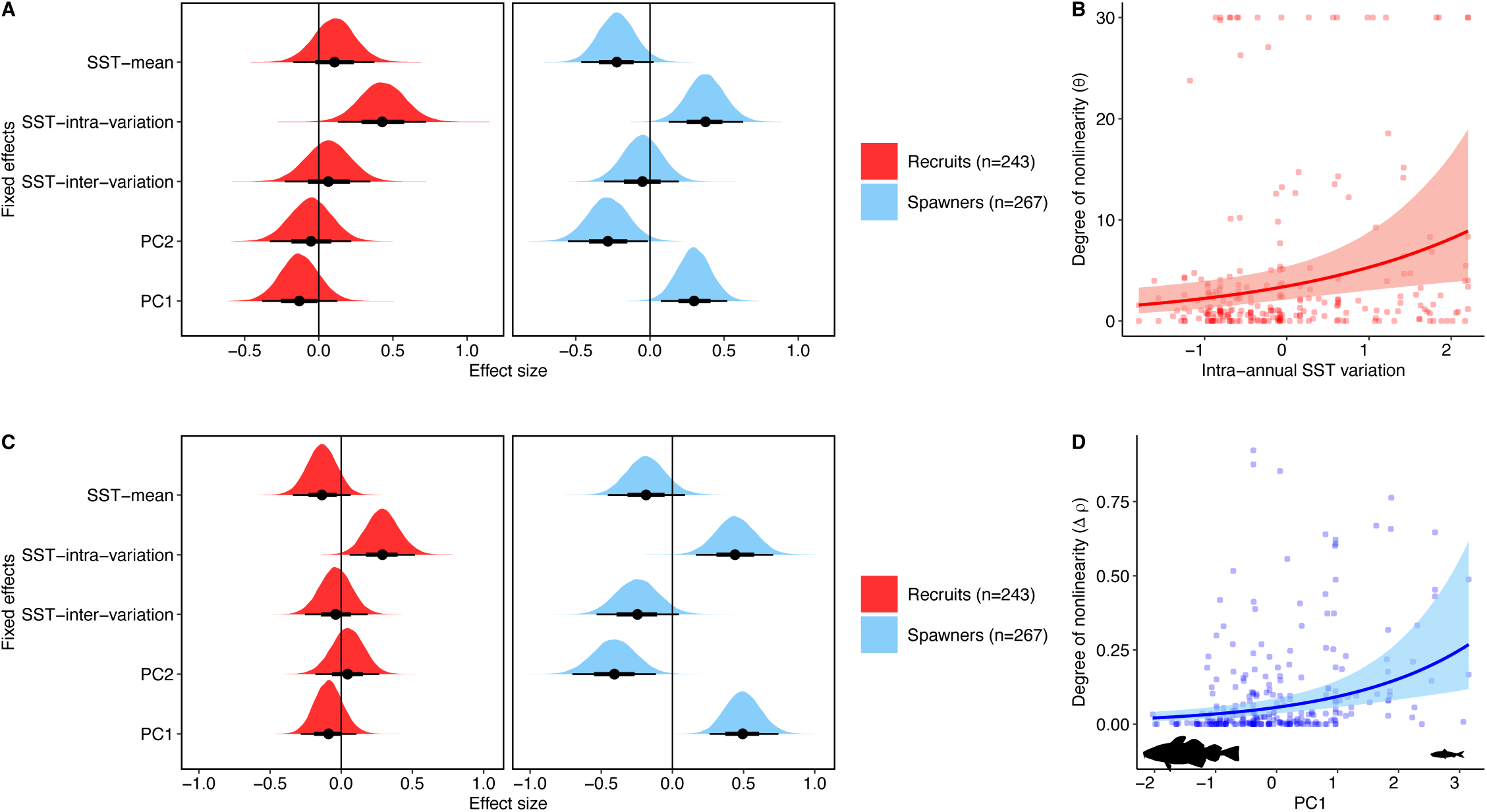
Temperature variation and fast life history traits amplify nonlinearity in marine fishes. Bayesian hurdle gamma models show consistent effects of sea surface temperature (SST) and life history correlates on two metrics of the degree of nonlinearity: **(A)** θ represents the S-maps nonlinear tuning parameter and **(C)** Δρ represents the difference in prediction accuracy between nonlinear and linear models. We present the posterior distributions along with a point estimate for the posterior median and the 66% (thick black line) and 95% (thin black line) credible intervals. All fixed effects were scaled to a mean of 0 and standard deviation of 1. Conditional effect plots show the degree of nonlinearity increasing with greater **(B)** intra-annual SST variation and **(D)** PC1 values representing traits of species with fast life-histories such as small body size (see fig. S2. for principal component analysis of life history traits). Line and shaded areas represent the median and 95% credible interval. Red and blue dots represent the values of θ and Δρ for 243 recruit and 267 spawner populations, respectively.

**Table 1.**
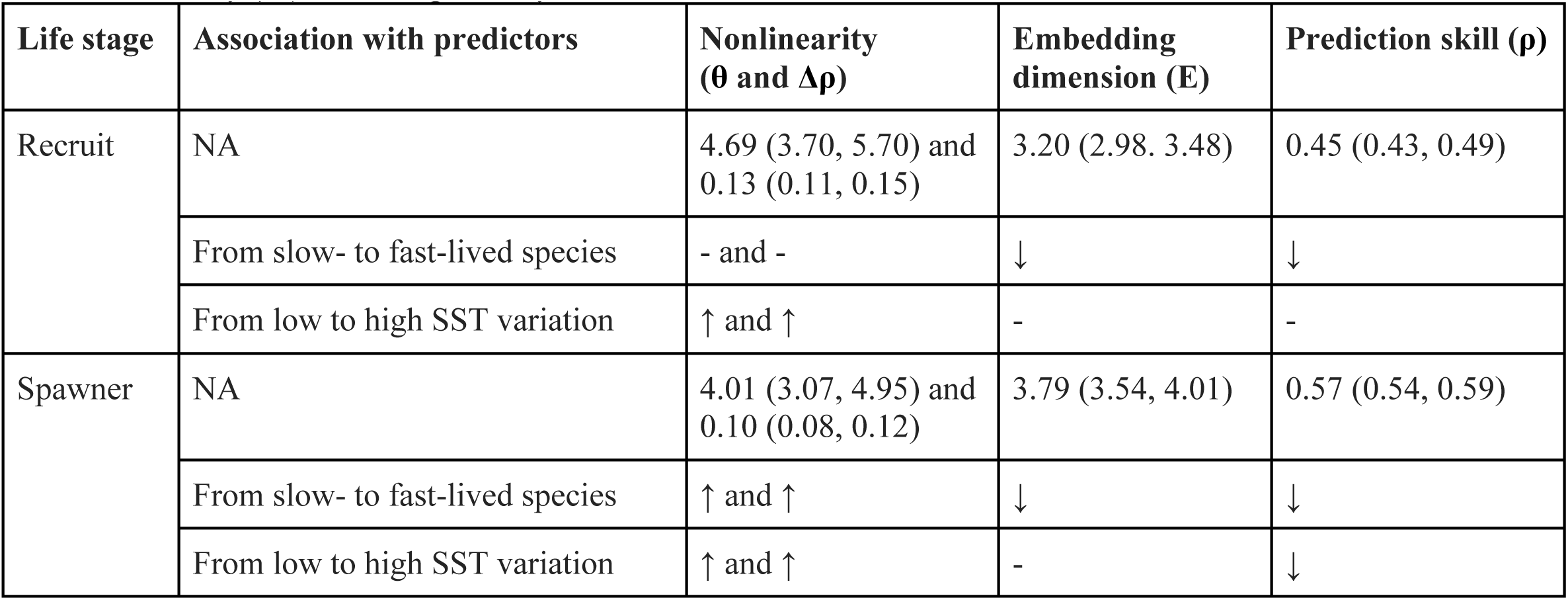
Summary of the main results. Mean values and 95% credible intervals in brackets of the empirical dynamic modeling metrics estimated by linear regressions. Associations between the metrics and life history traits and intra-annual sea surface temperature (SST) variation were evaluated using generalized linear mixed models. Positive, negative and no association are indicated by ↑, ↓ and -, respectively.

There was no clear difference in the mean values of θ and Δρ of recruitment and spawner biomass as indicated by largely overlapping 95% CI (Table 1). The ranges of θ and Δρ were 0 to 30 and 0 to 0.92, respectively (fig. S8). We found positive correlations between the coefficient of variation (CV) in time series and the degree of nonlinearity (figs. S9-10). The correlation between CV in recruitment and spawner biomass time series, respectively, was stronger for Δρ (95% CIs = -0.03 to 0.55 and 0.01 to 0.36) than θ (95% CIs = -0.001 to 0.01 and -0.0002 to 0.008; see fig. S11 and Supporting Text for further details). We also tested for a temporal trend in nonlinearity for the populations with 60+ time points and found that θ and Δρ values were greater in the second half of the time series compared to the first half for 31 (47%) and 34 (52%) of the 66 recruitment time series, respectively, and for 44 (46%) and 41 (43%) of the 96 spawner time series.

Embedding dimension (E) ranged from 1 to 10, and was on average greater for spawner biomass than recruitment time series (Fig. 1B, Table 1). Recruitment E increased with PC2, representing high maximum age and late age at maturity (95% CI = -0.03 to 0.67), and was not associated with the other predictors (Fig. 3A). Spawner biomass E increased with negative values of PC1, representing large body size and high trophic level (95% CI = -0.53 to -0.01). Spawner biomass E was weakly associated with PC2 (95% CI = -0.06 to 0.50), mean SST (95% CI = -0.07 to 0.48), intra-annual SST variation (95% CI = -0.47 to 0.11) and inter-annual SST variation (95% CI = -0.10 to 0.54).

**Fig. 3.**
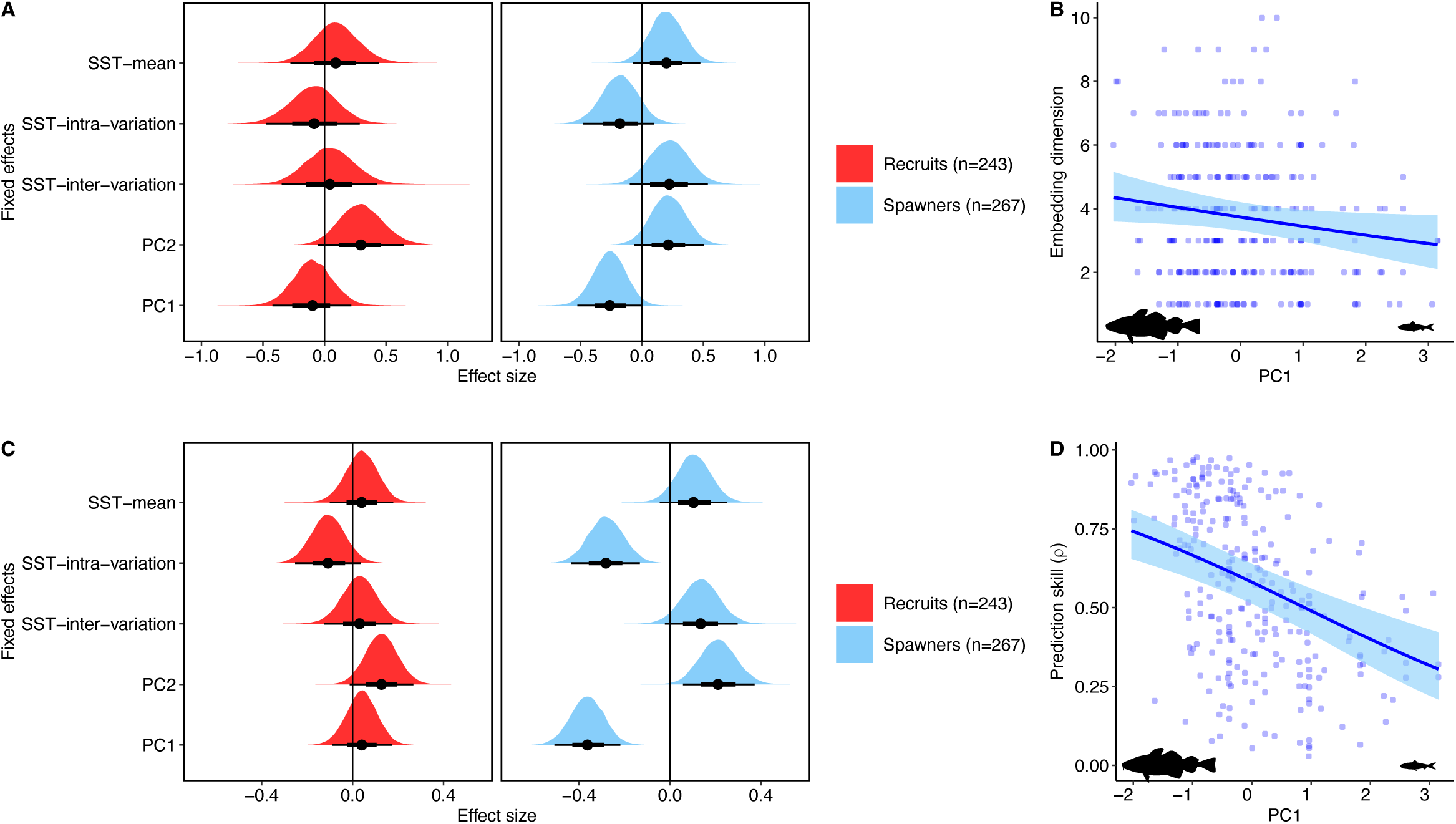
Embedding dimension and prediction skill are greater for species with slow life history traits. Bayesian cumulative and beta models show the effects of sea surface temperature (SST) and life history correlates on embedding dimension **(A)** and prediction skill **(C)**, respectively. We present the posterior distributions along with a point estimate for the posterior median and the 66% (thick black line) and 95% (thin black line) credible intervals. All fixed effects were scaled to a mean of 0 and standard deviation of 1. Conditional effect plots show **(B)** embedding dimension increasing with PC1 representing traits of slow-lived species and **(D)** prediction skill decreasing with increasing intra-annual SST variation. Line and shaded areas represent the median and 95% credible interval and blue dots represent the values of 267 spawner populations.

Prediction skill (ρ) ranged from 0.01 to 0.97, and was on average greater for spawners than recruitment (Table 1), and increased for species with slow life histories (Fig. 3C). Recruitment ρ increased with PC2 (95% CI = -0.01 to 0.27), representing high maximum age and age at maturity, and decreased with intra-annual SST variation (95% CI = -0.25 to 0.04). Recruitment ρ was not associated with the remaining predictors (Fig. 3C). Spawner biomass ρ increased with PC2 (95% CI = 0.06 to 0.37) and decreased with PC1 (95% CI = -0.51 to -0.22) and intra-annual SST variation (95% CI = -0.44 to -0.13). Spawner biomass ρ was weakly associated with mean SST (95% CI = -0.04 to 0.25) and inter-annual SST variation (95% CI = -0.03 to 0.29).

We further investigated the importance of SST on fish dynamics using convergent cross mapping (CCM), which identifies causation by detecting signatures of causal variables within the time series of affected variables^33^. We used Canada’s Northern Atlantic cod to demonstrate CCM (Fig. 4A) and to clarify longstanding debates regarding the influence of climate on this population (e.g. ^34,35^). The results show causal forcing of SST on cod as the attractor constructed from lags of the cod timeseries predicted SST (terminal library size ρ > 0.1, t-test p-value < 0.05) and prediction skill improved as the attractor was filled out with more data points (Kendall’s τ > 0, p-value < 0.05). The reverse is not possible as cod intuitively do not influence SST. We identified causal forcing of SST on 163 (67%) recruitment, 144 (54%) spawner biomass, and 196 (63%) total catch time series (Fig. 4B). The probability that recruitment and spawner biomass were causally forced by SST increased with fast-lived traits (Fig. 4C). For recruitment, the probability of causal SST forcing increased with intra-annual SST variation (95% CI = -0.01 to 1.13) and negative PC2 values (95% CI = -1.25 to -0.10) indicating fast intrinsic growth rates and high fecundity. For spawner biomass, the probability of causal SST forcing increased with positive PC1 values (95% CI = -0.03 to 0.74) representing fast somatic growth and young age at maturity. The probability that total catch was causally forced by SST increased for species more strongly associated with reef habitats (95% CI = -0.05 to 0.76) and was unaffected by other correlates.

**Fig. 4.**
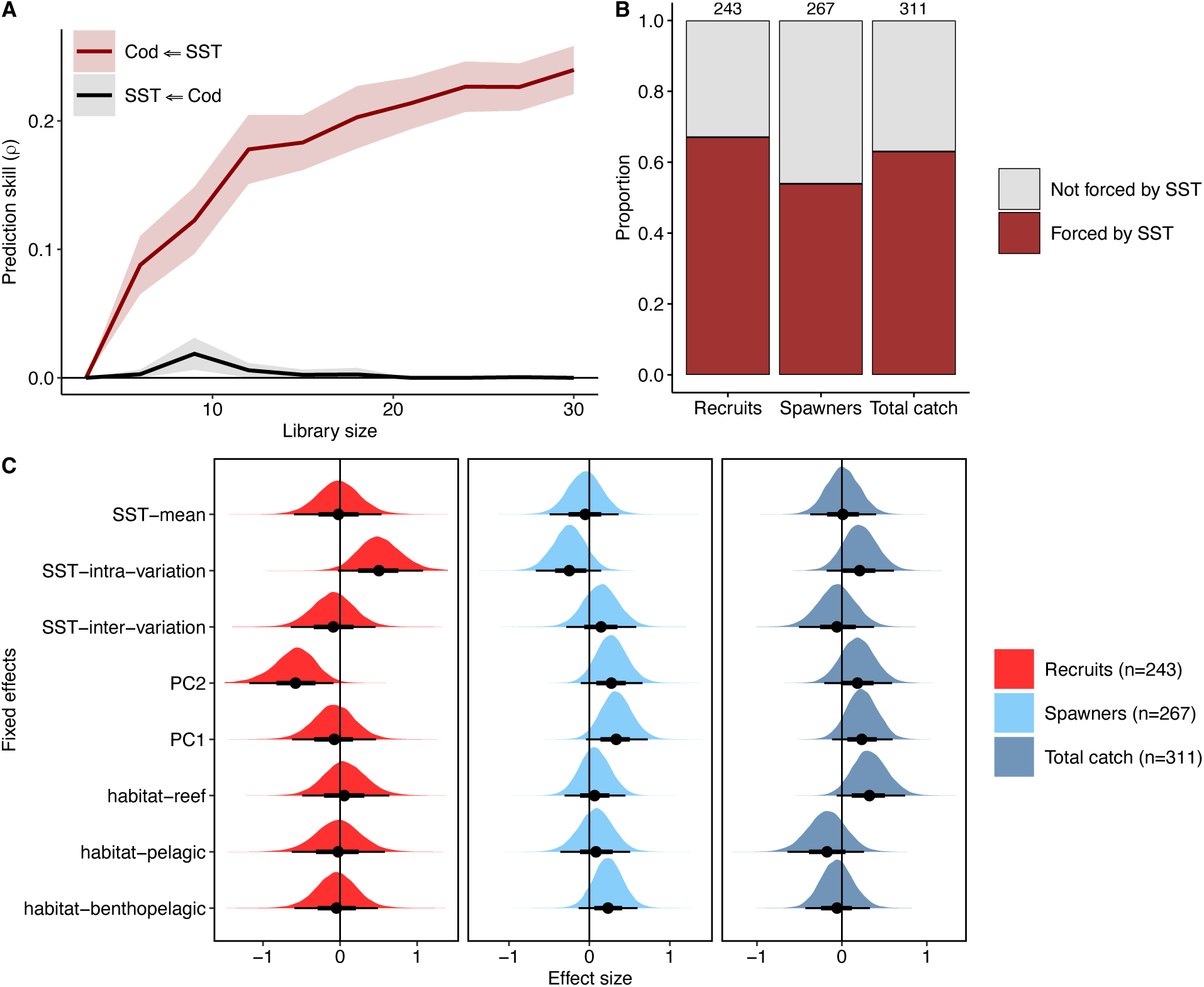
Widespread causal forcing of temperature on fish population dynamics. **(A)** Convergent cross mapping (CCM) between time series of Canadian Northern Atlantic Cod recruits and SST of the population’s boundary demonstrates unidirectional causal forcing of SST on cod. We show that the mean prediction skill (dark red line and shaded 95% confidence interval) increases as more data points are added to the cod attractor, which is possible because causal SST forcing has embedded a signature into the timeseries of cod. Legend arrows indicate forcing direction. **(B)** Proportion of global populations causally forced by SST as identified by CCM. Values above the bars represent the number of time series analyzed. **(C)** Bayesian Bernoulli models show the effects of SST, life history and habitat correlates on the probability that a population is causally forced by SST. We present the posterior distributions along with a point estimate for the posterior median and the 66% (thick black line) and 95% (thin black line) credible intervals. All fixed effects were scaled to a mean of 0 and standard deviation of 1.

## Discussion

Our results demonstrated that nonlinear fluctuations are common in populations of marine fish worldwide and are driven by both environmental and life-history factors, depending on life stage. Nonlinearity was amplified by intra-annual SST variation for both recruitment and spawner biomass, while life history mediated the degree of nonlinearity for spawners, dampening it in slow-lived species. We found fish population dynamics to collapse onto low-dimensional attractors. The mean embedding dimension was lower for recruits than spawners, indicating more rapid recurrence times and fewer processes underlying the dynamics of recruits compared to spawners. Prediction skill decreased for species with fast life history traits, suggesting that rapid and highly nonlinear dynamics decrease forecast accuracy. These nonlinear, low-dimensional fish population dynamics were often causally forced by SST, with a greater probability of casual forcing for recruits in variable SST environments and fast-lived recruits and spawners.

Nonlinearity in recruitment and spawner biomass time series increased with greater intra-annual SST variations (Fig. 2B), supporting the hypothesis that stochastic forcing is amplified by its interaction with nonlinearities^16^. Previous analyses, conditional on assumed parametric models, showed that elevated nonlinearity in fish is due to the amplification of process noise^2,8,24^, and demographic noise for *Daphnia*^25^. In our study, we used SST, and the results indicate that intra-annual variation provides the random variability that is amplified by the nonlinear behaviour of fishes. The deterministic source of nonlinearity may involve damped oscillations, as predicted by consumer-resource and food web theory^36^, observed in trophic cascades in marine ecosystems^37^, and predicted by the behaviour of simple fishery spawner-recruit data fitted to stock assessment time series^2,24^ and experimental manipulations of *Daphnia* populations^25^. Our study indicates that fish population fluctuations will become more nonlinear as a result of increasing environmental variability associated with global change^11^. This conclusion is further supported by ∼46% of analyzed time series already showing higher nonlinearity in the latter half of their time series compared to the beginning.

We found life history traits to mediate the degree of nonlinearity in spawner biomass (Fig. 2). It is well appreciated that populations of slow-lived species consist of more age classes that can smooth out environmental effects via portfolio effects and bet-hedging strategies^10,38,39^. As a result, species with slow life history traits dampened nonlinearity in spawner biomass because the variability is averaged across more age classes. On the other hand, fast-lived spawners displayed greater nonlinearity than those with slow life histories, which may reflect less buffering of recruitment nonlinearity. Our life history result also accords with the knowledge that fishing increases the degree of nonlinearity in marine fishes^8^. This stems from fishing-induced age-truncation, which reduces average body size and increases the intrinsic growth rate of fished compared to unfished populations^8,15^. Our study therefore supports the hypothesis that fishing-induced age-truncation increases fluctuations in fish abundance due to elevated nonlinearity^8,15^.

Warmer waters may be expected to increase nonlinearity due to elevated vital rates for ectothermic fish^40^, but we found the opposite effect for spawner biomass (Fig. 2). One potential explanation could be that nonlinear signatures are more apparent in colder environments due to less complex food webs^41^. We also found negative and positive effects of inter-and intra-annual SST variation on the nonlinearity of spawner biomass (Fig. 2). The former suggests there is a mechanism with a low degree of nonlinearity that enables spawners to withstand the lower mean inter-(∼0.3°C) than intra- (∼8°C) annual SST variation. Whereas the positive effect of the latter indicates that noise from within year temperature variation sustains the nonlinearity of spawner biomass that originated during the recruitment process.

The lower embedding dimension (E) of recruits than spawners (Fig. 1B) indicates that the dynamics of the former are faster than the latter as Eτ approximates the recurrence time - the time needed to return to a nearby state^30^. E also provides a heuristic measure of complexity, though it does not directly reflect the native dimensionality of the system (i.e. true number of state variables). Specifically, E is the minimum number of time-delayed coordinates needed to reconstruct the attractor for accurate prediction. In this sense, E represents the processes interacting to generate the observed dynamics and can be practically useful for informing the number of variables required to build a mechanistic model. The results (Figs. 1B and 3A) indicate that only ∼3-4 variables are needed to predict fish population fluctuations. This reinforces the low-dimensionality of nonlinear dynamics^42^ compared to high-dimensional stochastic systems (E>20)^9^, and is further supported by the negative correlations between E and nonlinearity metrics (Table S1). However, E is greater for spawners than recruits, and increased with slow-lived traits (Figs. 3A and 3B) suggesting that such life histories allow for more interactions with abiotic and biotic variables, though longer internal time lags and stochasticity could also contribute^9^. Our result contrasts previous work showing E to increase for animals with fast life history traits^6^. However, this discrepancy can be explained by added dimensionality from human intervention, as fisheries which target older, larger fish, are influenced by regulation and market prices^18,43,44^.

We detected widespread causal forcing of SST on fish dynamics (Fig. 4B). That a larger proportion of recruits were causally forced by SST than spawners (Fig. 4B), and that greater intra-annual SST variation increased the probability of forcing for recruits but not spawners (Fig. 4C), suggests that SST may be relatively more important for mediating nonlinearity in recruits than spawners (Fig. 2). While there was no association with SST summary metrics for spawners, fast life history traits increased the probability of SST forcing for spawners and recruits (Fig. 4C). These results are likely because temperature is critical in determining the growth of small, young fish, as well as food and habitat availability, whereas larger fish can smooth out environmental effects, as discussed previously^10,38,45^. We also detected causal SST forcing on ∼60% of global fisheries, which could reflect SST influencing population dynamics, as demonstrated above, and/or fishing effort and economic factors by determining the distribution of fish^46^. The higher probability of SST forcing on reef-associated fisheries may reflect temperature mediated dynamics of reefs, such as bleaching, as they provide habitat for many fishes^47,48^.

Conventional models for fisheries management assume a stable equilibrium exists, which is the basis for maximum sustainable yield, with random fluctuations caused by the environment^49^. However, our results support the view that nonlinear dynamics are ubiquitous^6^, which has implications for forecasting. For instance, prediction skill increased for slow-lived species (Fig. 3D) which may be possible because they do not linearly track stochastic changes in the environment. Rather, they display a low degree of deterministic nonlinearity that can be harnessed for accurate short-term prediction^21,22,42^ (Figs. 1C and 2C). In contrast, the high nonlinearity in fast-lived species may make them more difficult to predict (Fig. 2C). This is supported by prediction skill decreasing with greater intra-annual SST variation (Fig. 3C) which amplified nonlinearity in the preceding analyses (Fig. 2). Factors that increase nonlinearity, such as greater variability from global change and fishing-induced age truncation, may therefore have the undesirable effect of decreasing predictability. The most concerning populations from a management perspective may be those exhibiting high nonlinearity and low predictability, as they are likely susceptible to the largest noise amplification and irregular fluctuations. Several populations with well-documented management difficulties and complex population dynamics met these criteria (Table S2). For example, Atlantic cod (COD3Pn4RS, CODBA2224), Atlantic herring (HERR4TFA, HERR4TSP) and Pacific herring (HERRPRD, HERRWCVANI) that are characterized by population oscillations, abrupt collapses and subsequent commercial fishing moratoria (e.g. refs. ^35,37,50,51^). Harnessing nonlinearity may offer a powerful path toward effective and adaptive fisheries management^21,52,53^.

A logical interpretation of our results is that environmental variability drives nonlinearity early in life which is then propagated to fluctuations in the adult population, where life history traits modulate the degree to which it is expressed. This view is supported by nonlinear dynamics arising in marine fish via density-dependence during recruitment processes as young individuals compete for resources and are subject to intense predation^20,54–56^. That nonlinearity arises through recruitment accords with conventional knowledge in fisheries stock assessment, where density dependence is often represented in recruitment for spawner-recruit relationships^49,56^.

Accordingly, most recruitment time series displayed nonlinear dynamics (Fig. 1C) and were causally forced by temperature (Fig. 4B). Recruitment nonlinearity was amplified by temperature variation (Fig. 2) and the probability of the dynamics being causally forced by temperature increased for recruits with greater temperature variation (Fig. 4C). Spawner nonlinearity increased with both temperature variation and fast-lived traits (Fig. 2), and the probability of causal temperature forcing increased for fast-lived species (Fig. 4C). These results are consistent with younger individuals and fast-lived species being more temperature sensitive while slow-lived species buffer environmental variability through age-class integration and bet-hedging^10,38,39,45^. Our view is therefore that temperature variability amplifies recruitment nonlinearity, which is propagated to spawners and further amplified by temperature in fast-lived species but dampened in slow-lived species. A causal demonstration of this pathway is warranted and presents an intriguing avenue for future research.

Overall, our study demonstrated that nonlinear dynamics are widespread in populations of exploited fish, correlated with the magnitude of fluctuations in time series, and driven by both intrinsic and extrinsic factors. The results indicate that temperature variation amplifies nonlinearity in recruitment, while life-history traits may mediate the degree to which nonlinearity is propagated to spawner biomass, dampening it in species with slow life histories. We found fish dynamics to collapse onto low-dimensional attractors. The embedding dimension was greater for spawners than recruits, and increased with slow life history traits indicating that fishing of larger, older spawners increases complexity. Causal forcing of SST on fish dynamics was more pronounced in recruitment than spawner time series, and the probability of forcing increased for recruits in variable temperature environments and spawners with fast life histories, consistent with our nonlinearity findings. Collectively, our study indicates that marine fish populations may experience higher amplitude and/or more frequent boom-and-busts that are difficult to predict as a result of increased nonlinearity from fishing-induced age-truncation and elevated environmental variability associated with global change.

## Author contributions

Conceptualization: RMH, MK. Methodology: RMH, MK. Formal analysis: RMH. Visualization: RMH. Supervision: MK. Writing – original draft: RMH. Writing – review & editing: RMH, MK.

## Acknowledgments

We are grateful to the many individuals and organizations that contributed to the databases that made our study possible.

## Data and materials availability

The RAM Legacy Stock Assessment Database^26^ and NOAA Optimum Interpolation SST V2 High Resolution database^28^ are publicly available. We have archived all code and analyzed data on a repository which will be made publicly available upon manuscript acceptance.

## Materials and methods

### Data collection

We extracted annual spawner stock biomass, recruitment and total catch time series of non-salmonid, iteroparous marine fish species from the RAM Legacy Stock Assessment Database, hereafter referred to as RAM LSAD^26^. The time series of spawners represents the biomass of reproductively mature adults, while recruitment is the number of young fish produced by spawners that survive early density-dependent mortality and subsequently ’recruit’ into the population. Our main nonlinearity analysis focused on populations that had at least 30 continuous years of data between 1982 and 2022, aligning with the available sea surface temperature (SST) data from the same period. We excluded early values in the time series that were not strongly influenced by data, following the methods of a previous analysis of the RAM LSAD^29^, which included identifying and removing initial periods of flat or smoothly increasing trends in recruitment or spawner biomass. This resulted in 273 spawner and 248 recruitment time series. Six spawner and five recruit populations had negative prediction skills across all tested values of θ (see Empirical Dynamic Modeling section below) and were therefore excluded from further analysis. As a result, our final dataset for the main analysis included time series of 267 spawners and 243 recruit populations, covering 141 spawner species and 131 recruit species, for a total of 143 species (Fig. S1). To visualize the geographic extent of this dataset, we plotted the closest ocean cell to the mean latitude and longitude of each fish population, using the population boundaries previously delineated^29^ (Fig. 1). The mean time series length was 56.2 years for spawners (standard deviation = 24.1) and 51.3 for recruitment (standard deviation = 18.3).

For our main analysis, we extracted SST time series spanning 1982 to 2022 from the NOAA Optimum Interpolation SST V2 High Resolution database due to its 0.25 by 0.25 degree fine resolution^28^. For our fourth sensitivity analysis (see section below), we extracted SST time series from the COBE-SST 2 database which is more coarse (1 by 1 degree)^58^. From both SST databases, we used stock boundaries delineated^29^ to create mean annual SST time series for each population. Intra-annual SST variation for each population was calculated by averaging the difference between the maximum and minimum monthly SST within a year. Inter-annual SST variation was calculated by averaging the absolute first differences of the annual mean SST time series.

We used FishLife to estimate life history traits at the species level when possible, or genus level when species-specific data was not available^27,59^. For example, the RAM LSAD lists the scientific name for sand lance stocks, such as SEELVIa, as Ammodytes spp. We performed principal component analyses of the following life history traits: intrinsic growth rate (r), trophic level, log-max length, log-max age, log-age at maturity, log-fecundity, log-growth coefficient (Fig. S2). We also extracted trait values for habitat associations from FishLife: habitat-reef-associated, habitat-pelagic and habitat-benthopelagic.

### Empirical dynamic modeling

We used empirical dynamic modeling (EDM), a data-driven framework, for the nonlinear time series analyses^30^, and performed all analyses with leave-one-out cross-validation in R using the rEDM package (version 1.15.4)^60^. Following best practices of EDM, we first differenced and scaled all time series to mean = 0 and standard deviation = 1. We used simplex projection via the EmbedDimension function to determine the embedding dimension, E, which is the number of time lags needed to reconstruct the system in the state-space. We tested values between 1 and the square root of the time series length, up to a maximum of 10 and selected the value of E that maximized out-of-sample prediction skill (ρ) via leave-one-out cross-validation. We used sequentially weighted global linear maps (S-maps) to quantify nonlinear dynamics in spawner and recruitment time series, using the optimal E and where the nonlinear tuning parameter, θ, controls how much weighting is given to nearby points on the attractor^32^. At θ = 0, the S-map is a global linear model that weights all points on the attractor equally, but as θ > 0 the model becomes local nonlinear and more weight is given to points nearby on the attractor. We used rEDM’s PredictNonlinear function to test θ values from 0 to 30 at the hundredth decimal place and identified the optimal model as θ with the greatest out-of-sample ρ via leave-one-out cross-validation (Fig. S8). For S-map, we set the exclusion radius equal to E. For both simplex projection and S-map, we tested 𝜏 values of -1 and -2, and proceeded with -1 due to increased prediction skill. We used the following EDM parameters in our Bayesian models (see section below): two metrics of nonlinearity, 𝜃_ρ*max*_ and 𝛥𝜌 = 𝜌_*max*_ − 𝜌_θ=0_, E and ρ^2,7–9,16^. We performed five sensitivity analyses to account for uncertainty in the θ value, SST database and phylogenetic non-independence and found consistent results (see sensitivity analysis section).

We performed convergent cross mapping (CCM) between SST time series from the NOAA Optimum Interpolation SST V2 High Resolution database and the 267 spawner and 243 recruitment populations from the main nonlinearity analysis as well as 311 time series of total catch across 163 species from RAM LSAD. We used only data points between 1982 and 2022 for the fish time series to match the time frame of the SST database. This resulted in mean time series length and standard deviation, respectively, of 35.98 and 3.05 for recruitment, 36.61 and 3.05 for spawner biomass, and 35.87 and 3.06 for total catch. Following best practices of CCM^33^, we first differenced and scaled time series to mean = 0 and standard deviation = 1. We identified the optimal E (between 1 and square root of time series length) for each population that maximized ρ via leave-one-out cross-validation and used the maximum possible library size to compute CCM with random sampling, beginning at the smaller of 3 or E, and ending at the time series length – E + 2. We increased the library size in increments of 3, drew 100 random samples at each library size evaluated and set the exclusion radius equal to E. All CCM correlations were truncated at 0. We examined causation over time points -4 to 0 as there may be a time delayed causal effect of temperatures on system dynamics^61^. Following a one-sided t-test and Kendall’s τ test, we considered a population to be causally forced by SST if the following criteria were met for any of the time points: terminal ρ > 0.1 and t-test p-value <0.05, and if ρ increased with library size as indicated by Kendall’s τ > 0 and p-value < 0.05.

### Bayesian modeling

To analyze variation between recruitment and spawner time series in the outputs of the EDM analysis, we used Bayesian linear models to model θ, Δρ, E and ρ as functions of time series type (either spawner or recruitment). To analyze correlations between the coefficients of variation (CV) of the time series and nonlinearity, we modeled the CV of recruitment and spawner biomass as functions of θ and Δρ. All models assumed a gaussian distribution. We fit the models using the brms package^62^ in R^63^, with four MCMC chains that included 2000 warm-up iterations and 10000 sampling iterations. We also estimated associations between θ, Δρ, E and ρ via pairwise Bayesian correlation tests using the correlationBF function from the BayesFactor^64^ package in R.

We used Bayesian generalized linear mixed models (GLMMs) to analyze variation in θ, Δρ, E and ρ in relation to environmental and life-history correlates. We used hurdle gamma models for 𝜃_ρ*max*_ and 𝛥𝜌 = 𝜌_*max*_ − 𝜌_θ=0_ as both variables were positive, continuous, and included zero. We used cumulative logit models for E, as it is an ordinal categorical variable. We used binomial models for ρ, as the values were between 0 and 1. We used Bernoulli models to assess the probability of a population being influenced by SST, with the outcome being either forced or not forced. For the hurdle gamma, cumulative logit and binomial models we analyzed data from 267 spawner and 243 recruitment populations. For the Bernoulli models we also analyzed fisheries data from 311 total catch time series. For all GLMMs, we used the following fixed effects: mean SST, intra-annual SST variation, inter-annual SST variation and principal components 1 and 2 of a principal component analysis of intrinsic growth rate (r), trophic level, log-max length, log-max age, log-age at maturity, log-fecundity, and log-growth coefficient. For the Bernoulli models, we also used habitatreefassociated, habitatpelagic and habitatbenthopelagic as fixed effects. These represent the log-odds of association with the respective habitats (reef, pelagic, benthopelagic) relative to the baseline habitat, demersal. We scaled all fixed effects to mean 0 and a standard deviation of 1. We included species, stock area and assessment type as random effects in all models to account for variability across species, geographical locations and stock assessment type. We confirmed model convergence and fit based on trace plots, posterior predictive check plots, effective sample sizes greater than 1,000, and Gelman-Rubin convergence diagnostic values (R-hat) equal to 1^65^. We consider strong effects as those where the 95% credible interval excludes 0 and the posterior distribution is clearly shifted away from 0, weak effects where the 95% CI includes 0 but most of the posterior distribution is shifted away from 0, and no effect where the 95% CI includes 0 and the posterior distribution is not shifted away from 0.

### Sensitivity analyses

#### Sensitivity analysis 1 and 2

We performed multiple sensitivity analyses to account for uncertainty in the value of θ. In the analysis presented in the main text (Fig. 2A) we tested θ values from 0 to 30 at the hundredth decimal and identified the best θ for each time series as the one that maximized ρ. However, it is possible that there is uncertainty in the best estimate of θ. To incorporate this uncertainty, we fitted 100 hurdle gamma models wherein a new θ value was randomly sampled, based on weight, for each population in every model. The weight was determined by the scaled distribution of 𝑤 = (𝜌_*max*_ − 𝜌)/𝜌_*max*_ over the range of θ where 𝑤 < 0.02 and separately 𝑤 < 0.035, that is, values of 𝜃 whose 𝜌 were within 2% and 3.5% of 𝜌_*max*_ (Fig. S11). For the 2% threshold (sensitivity analysis 1), this included on average 262 and 229 possible values of θ to randomly sample from for each spawner and recruit population, respectively. For the 3.5% threshold (sensitivity analysis 2), this included on average 362 and 319 possible values of θ to sample from for each spawner and recruit population, respectively. We then randomly sampled θ based on the weighting and re-ran our hurdle gamma models as before with fixed effects: mean SST, intra-annual SST variation, inter-annual SST variation and PC1 and PC2, and random effects for species, stock area and assessment method. We again used four MCMC chains with 2000 warm-up iterations and 10000 sampling iterations and evaluated model fit using trace plots, posterior predictive check plots, effective sample sizes greater than 1,000, and Gelman-Rubin convergence diagnostic values (R-hat) equal to 1^65^.

#### Sensitivity analysis 3

We selected the value of θ for all populations that produced the second greatest value of ρ. We again used a hurdle gamma model as before with fixed effects: mean SST, intra-annual SST variation, inter-annual SST variation and PC1 and PC2, and random effects for species, stock area and assessment method. We again used four MCMC chains with 2000 warm-up iterations and 10000 sampling iterations and evaluated model fit using trace plots, posterior predictive check plots, effective sample sizes greater than 1,000, and Gelman-Rubin convergence diagnostic values (R-hat) equal to 1^65^.

#### Sensitivity analysis 4

We also ran the θ hurdle gamma model using temperature correlates calculated from the COBE-SST 2 database which has a longer time series starting in 1850 but is more coarse (1 by 1 degree) than the NOAA OI SST V2 (0.25 by 0.25 degree). Using COBE-SST 2, we calculated the mean, intra and inter-variation SST correlates using the years that matched the fish time series. There were four spawner and three recruit populations from our main analysis that had to be removed as the COBE-SST 2 did not have SST values available for their stock boundaries. For example, we excluded populations from the Strait of Georgia, located between eastern Vancouver Island and mainland British Columbia, because the COBE-SST 2 dataset does not include data for this area, whereas the NOAA OI SST V2 dataset does. Thus, our hurdle gamma models included data from 240 recruit and 264 spawner populations. We again used four MCMC chains with 2000 warm-up iterations and 10000 sampling iterations and evaluated model fit using trace plots, posterior predictive check plots, effective sample sizes greater than 1,000, and Gelman-Rubin convergence diagnostic values (R-hat) equal to 1^65^.

#### Sensitivity analysis 5

Our fifth sensitivity analysis tested whether the results of our θ and Δρ hurdle gamma models held after correcting for phylogenetic non-independence. We obtained a time-calibrated phylogeny of ray-finned fishes^66^ which included 112/132 recruit and 119/138 spawner species used in our main analysis and omitted the populations for unavailable species from further analysis. We then ran our θ and Δρ models with the same fixed and random effects as the models presented in the main text. However, we also incorporated a species-level random effect with the values correlated according to the phylogenetic correlation matrix derived from a phylogeny of ray-finned fishes. We again used four MCMC chains with 2000 warm-up iterations and 10000 sampling iterations and evaluated model fit using trace plots, posterior predictive check plots, effective sample sizes greater than 1,000, and Gelman-Rubin convergence diagnostic values (R-hat) equal to 1^65^.

### Supporting Text

#### Extended description of the coefficient of variation results

We found positive correlations between the coefficient of variation (CV) in time series and the degree of nonlinearity, consistent with ref. ^7^. However, effect sizes varied between the two metrics of nonlinearity. The correlation between CV in recruitment and spawner biomass time series, respectively, was strong with Δρ (95% CIs = -0.03 to 0.55 and 0.01 to 0.36; Fig. S9) and weak with θ (95% CIs = -0.001 to 0.01 and -0.0002 to 0.008; Fig. S10). This can be explained by greater uncertainty in the value of θ than Δρ. Following best practices of empirical dynamic modeling, we selected 𝜃_ρ*max*_ as the optimal value. However, multiple values of θ produce qualitatively similar ρ values, as shown in the example (Fig. S11), which represents a common pattern we observed across the time series. In this example, there are 219 values of θ within 3.5% of 𝜌_*max*_, spanning θ values 2.04 to 4.22. On the other hand, Δρ ranges between 0.32 to 0.33. To quantify the variability, we calculated the standard deviation of θ and Δρ values within the 3.5% threshold for each time series, and subsequently modeled the standard deviation values as a function of the metric (either θ or Δρ). We found that the standard deviation was greater for θ than Δρ in both time series types. For spawner biomass, the difference in standard deviation between θ and Δρ was 1.13 (95% CI = 0.95 to 1.31), and for recruitment the difference was 1.01 (95% CI = 0.83 to 1.18). We believe the greater variation in θ than Δρ explains the differences in effect sizes.

We also found that the association between CV and EDM predictability (ρ) was positive for recruits (0.17, 95% credible interval = 0.04 to 0.29) but not clear for spawners (−0.06, 95% CI = - 0.18 to 0.06). This may indicate that the nonlinear signal in recruitment can be harnessed for prediction, while there is no clear effect for spawners.

#### Extended description of sensitivity analyses

We found consistent results across all sensitivity analyses. These analyses confirmed our results presented in the main text - that nonlinearity of recruits and spawners increased with greater intra-annual SST variation and nonlinearity of spawners increased with positive values of principal component 1 representing traits of slow-lived species.

Sensitivity analysis 1 (Fig. S3) used 100 hurdle gamma models and a 2% threshold which resulted in 262 and 229 mean possible θ values to randomly sample from for each spawner and recruit population, respectively. We found that for recruits, all 95% credible intervals (CI) were positive and non-zero overlapping for intra-annual SST variation. We did not find any clear effect from the other predictors. For spawners, we found that the 95% CI were positive and non-zero overlapping in 100 of 100 models for PC1, 94 of 100 models for intra-annual SST variation and negative and non-zero overlapping for 40 of 100 models for mean SST. We did not find clear effects from the other predictors.

Sensitivity analysis 2 (Fig. S4) used 100 hurdle gamma models and a 3.5% threshold which resulted in 362 and 319 mean possible θ values to randomly sample from for each spawner and recruit population, respectively. We found the results to be qualitatively similar as sensitivity analysis 1 but with more 95% CI overlapping 0 which is expected given the additional θ values. We found that for recruits, the 95% CI for intra-annual SST variation were positive and non-zero overlapping for 100 of 100 models. We did not find any clear effect from the other predictors.

For spawners, we found that the 95% CI for PC1 were positive and non-zero overlapping for 76 of 100 models, 68 of 100 models for intra-annual SST variation and for mean SST they were negative and non-zero overlapping for 28 of 100 models. We did not find any clear effect from the other predictors.

Sensitivity analysis 3 (Fig. S5) used the value of θ that produced the second greatest value of ρ. For recruits, there was a positive association with intra-annual SST variation (95% CI = 0.09 to 0.73). The remaining fixed effects were unclear. For spawners, there were associations with PC1 (95% CI = 0.19 to 0.69), PC2 (95% CI = -0.55 to 0.04), intra-annual SST variation (95% CI = 0.06 to 0.61) and mean SST (95% CI = -0.56 to -0.03). The remaining fixed effects were unclear.

Our sensitivity analysis 4 (Fig. S6) used SST correlates generated from the COBE database. For recruits, nonlinearity was associated with intra-annual SST variation (95% CI = 0.17 to 0.73).

The remaining fixed effects were unclear. For spawners, nonlinearity was associated with PC1 (95% CI = 0.09 to 0.53), PC2 (95% CI = -0.53 to 0.01), intra-annual SST variation (95% CI = 0.11 to 0.58) and mean SST (95% CI = -0.47 to 0.02). The remaining fixed effects were unclear.

Sensitivity analysis 5 (Fig. S7) shows that the main results held after correcting for phylogenetic non-independence using a phylogenetic correlation matrix random effect. Importantly, the relationships remained consistent between nonlinearity metrics and intra-annual SST variation for recruits (95% CI for θ = -0.01 to 0.62 and Δρ = 0.00 to 0.50) and spawners (95% CI for θ = 0.11 to 0.65 and Δρ = 0.17 to 0.76) and PC1 for spawners (95% CI for θ = -0.11 to 0.40 and Δρ = -0.03 to 0.53). The reduced clarity in association with spawner PC1 for θ is likely due to the greater uncertainty in the value of θ than Δρ (fig. S6.B), as discussed above. Nonlinearity in spawner biomass was also associated with inter-annual SST variation (95% CI for θ = -0.50 to 0.08 and Δρ = -0.65 to -0.03).

**Table S1.** Pairwise associations between EDM metrics as estimated by Bayesian correlation tests. We show the median of the posterior distribution as well as the 95% credible intervals in brackets. Bold (top) and regular font (bottom) indicate correlations for recruitment and spawners, respectively.

| | $\theta$ | $\Delta\rho$ | E | $\rho$ |
| --- | --- | --- | --- | --- |
| $\theta$ | 1<br>1 | | | |
| $\Delta\rho$ | **0.35 (0.24, 0.47)**<br>0.58 (0.50, 0.66) | 1<br>1 | | |
| E | **-0.08 (-0.20, 0.05)**<br>-0.12 (-0.23, 0.00) | **-0.23 (-0.34, -0.10)**<br>-0.18 (-0.29, -0.07) | 1<br>1 |  |
| $\rho$ | **-0.15 (-0.28, -0.03)**<br>-0.46 (-0.55, -0.36) | **0.03 (-0.10, 0.15)**<br>-0.43 (-0.52, -0.33) | **-0.10 (-0.22, 0.02)**<br>0.08 (-0.04, 0.19) | 1<br>1 |

**Table S2.** The top 20 spawner populations ranked by high nonlinearity and low predictability. Ranking was determined by summing the ranks assigned in descending order of nonlinearity (θ) and ascending order of predictability (ρ). Population ID is the stockid from the RAM LSAD.

| Rank | Species | Population (stock) ID | $\theta$ | $\Delta\rho$ | $\rho$ |
| --- | --- | --- | --- | --- | --- |
| 1 | *Clupea harengus* | HERRRIGA | 30.00 | 0.05 | 0.06 |
| 2 | *Lepidorhombus whiffiagonis* | MEG8c9a | 30.00 | 0.41 | 0.09 |
| 3 | *Clupea harengus* | HERR4TFA | 28.84 | 0.49 | 0.08 |
| 4 | *Sebastes nebulosus* | CHROCKNP coast | 26.89 | 0.06 | 0.08 |
| 5 | *Gadus morhua* | COD3Pn4RS | 30.00 | 0.92 | 0.15 |
| 6 | *Clupea harengus* | HERR4TSP | 18.29 | 0.08 | 0.05 |
| 7 | *Clupea pallasii* | HERRPRD | 16.74 | 0.32 | 0.07 |
| 8 | *Dissostichus eleginoides* | PTOOTHFISHPEI | 27.72 | 0.23 | 0.14 |
| 9 | *Pleuronectes platessa* | PLAICIS | 20.10 | 0.13 | 0.13 |
| 10 | *Solea solea* | SOLEIIIa-2224 | 12.14 | 0.26 | 0.07 |
| 11 | *Thunnus alalunga* | ALBAIO | 11.83 | 0.15 | 0.09 |
| 12 | *Trachurus japonicus* | JMACKPJPN | 8.71 | 0.60 | 0.03 |
| 13 | *Gadus morhua* | CODBA2224 | 20.79 | 0.41 | 0.18 |
| 14 | *Clupea harengus* | HERRNIRS | 21.70 | 0.61 | 0.25 |
| 15 | *Engraulis encrasicolus* | ANCHOBAYB | 19.49 | 0.65 | 0.28 |
| 16 | *Melanogrammus aeglefinus* | HAD4X5Y | 22.18 | 0.20 | 0.29 |
| 17 | *Mallotus villosus* | CAPENOR | 30.00 | 0.66 | 0.32 |
| 18 | *Brevoortia patronus* | MENATGM | 6.69 | 0.06 | 0.14 |
| 19 | *Engraulis encrasicolus* | ANCHMEDGSA29 | 30.00 | 0.45 | 0.33 |
| 20 | *Clupea pallasii* | HERRWCVANI | 29.56 | 0.18 | 0.32 |

**Fig. S1.**
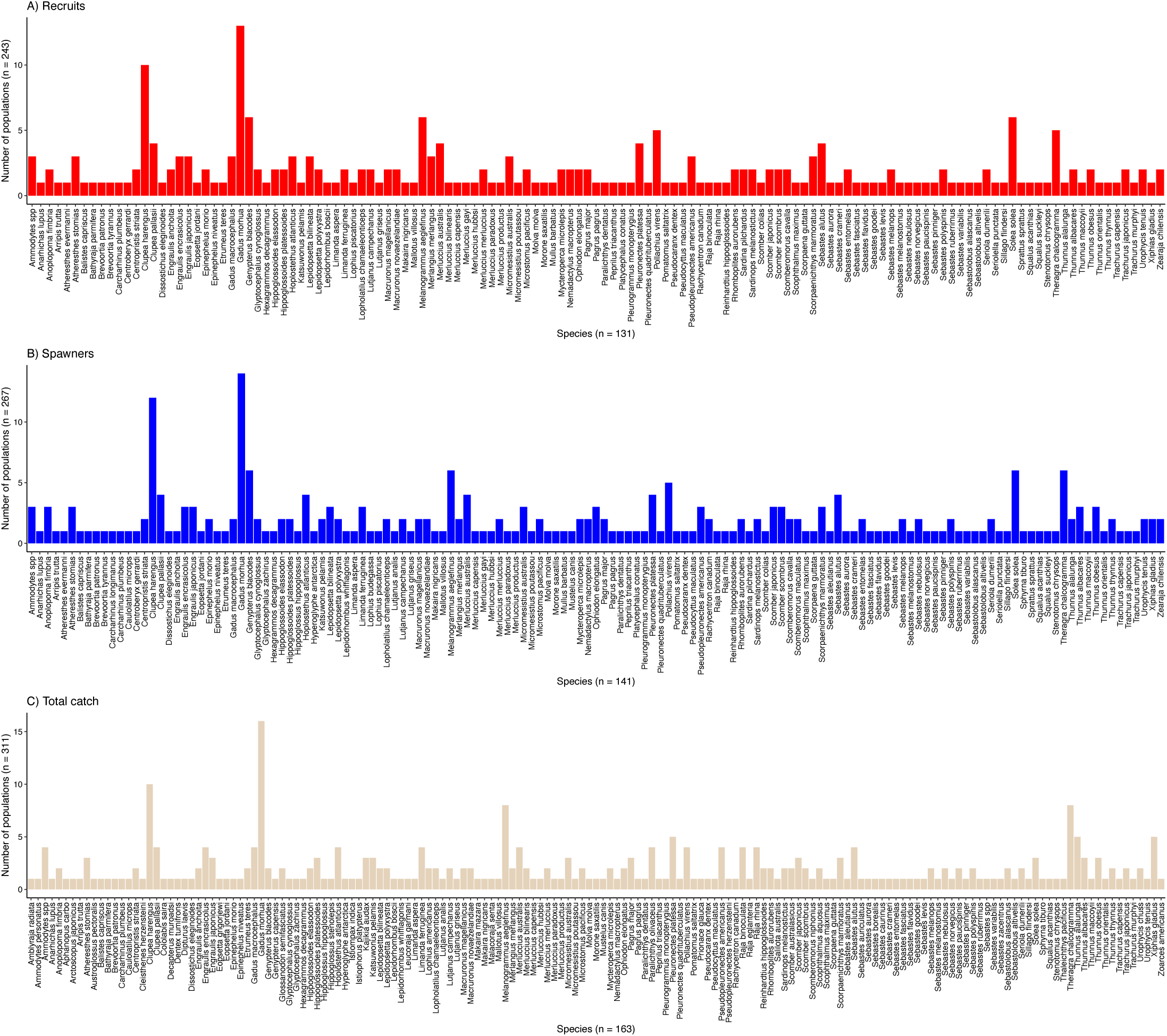
Histograms showing the number of populations per species used in our analyses.

**Fig. S2.**
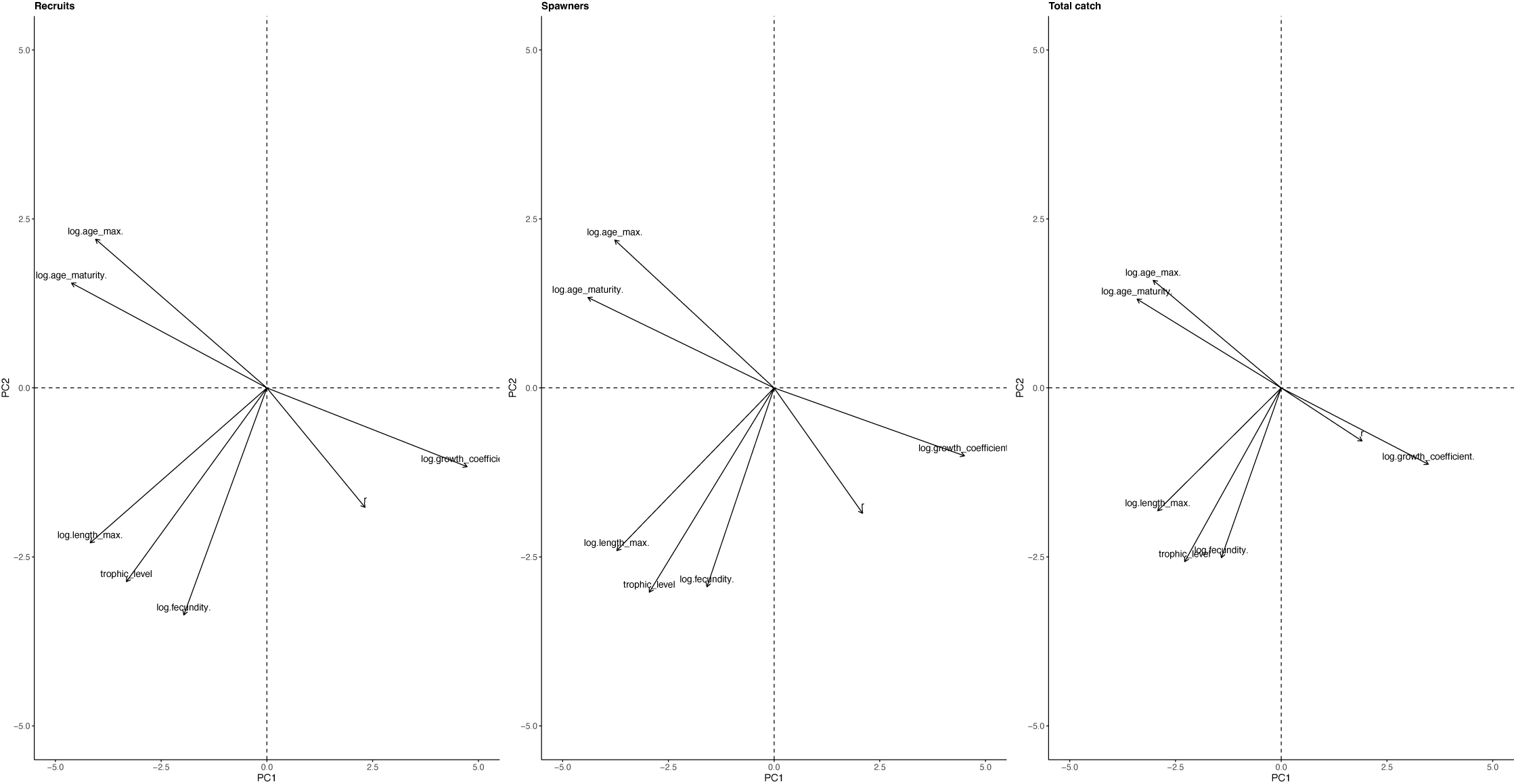
Principal component analyses of species life history traits: intrinsic growth rate (r), trophic level, log-max length, log-max age, log-age at maturity, log-fecundity and log-growth coefficient. We used PC1 and PC2 as predictors for our models.

**Fig. S3.**
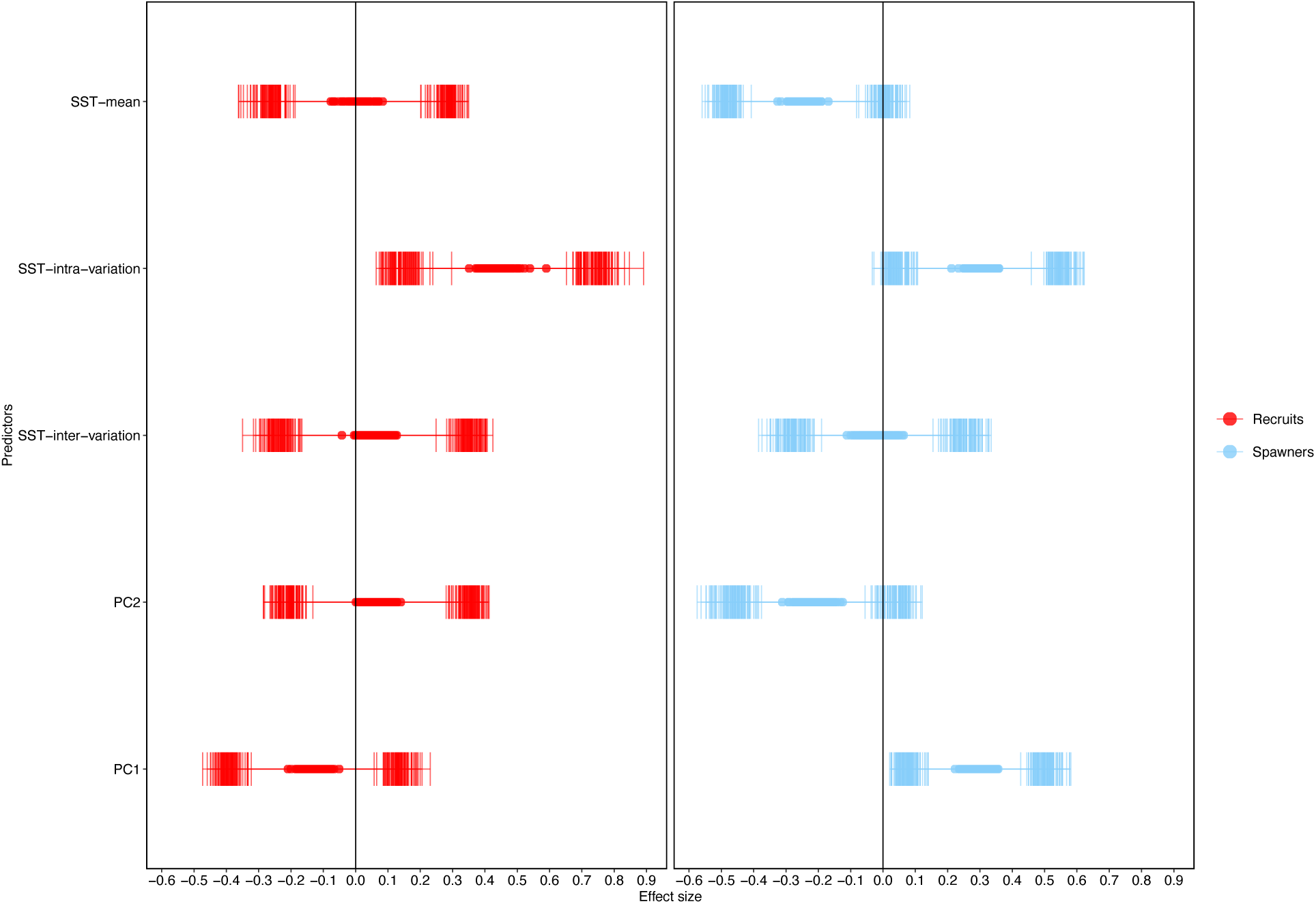
Sensitivity analysis 1 using the 2% threshold. We show the median point estimates and 66% and 95% credible intervals of our predictors for 100 hurdle gamma models.

**Fig. S4.**
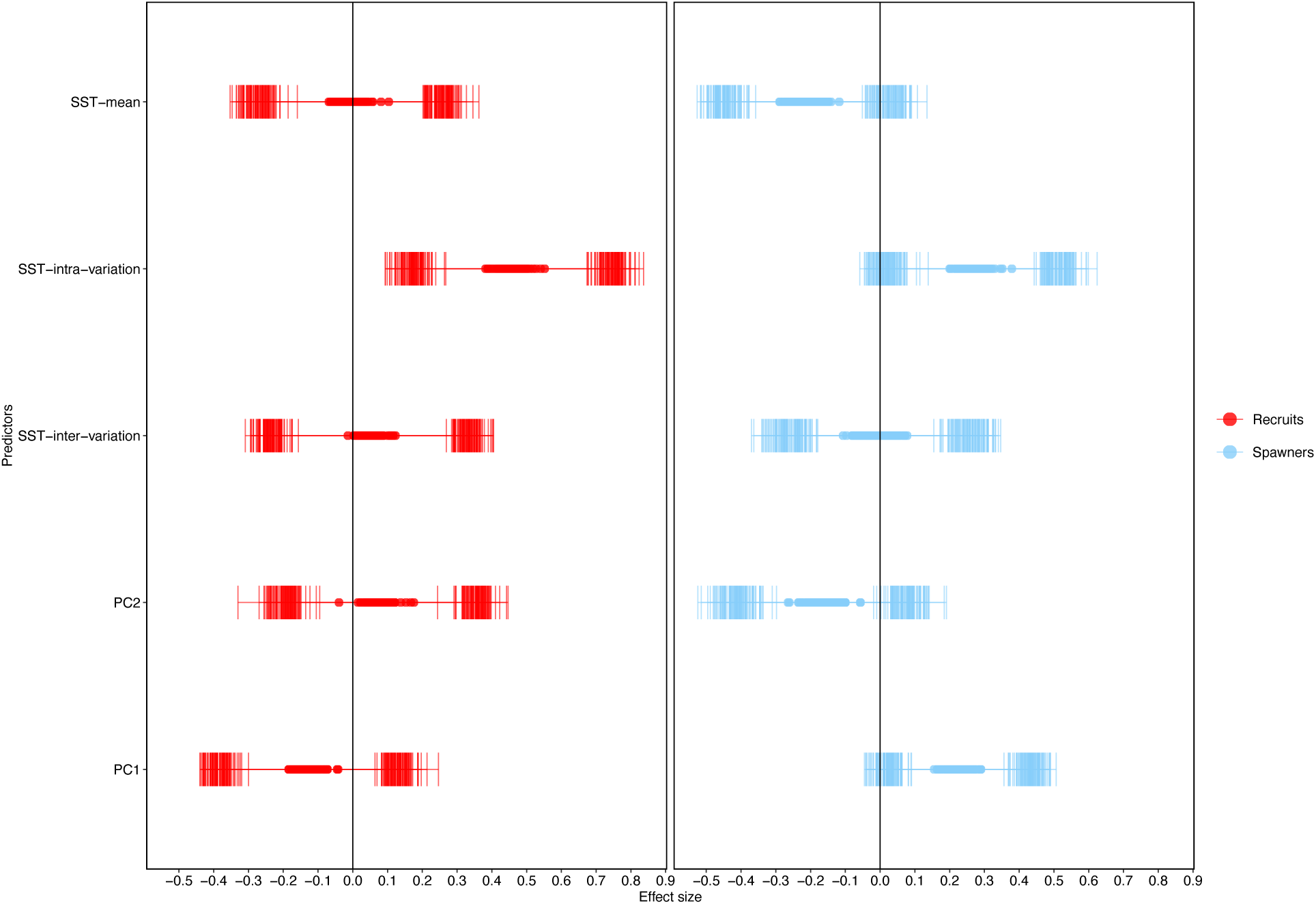
Sensitivity analysis 2 using the 3.5% threshold. We show the median point estimates and 66% and 95% credible intervals of our predictors for 100 hurdle gamma models.

**Fig. S5.**
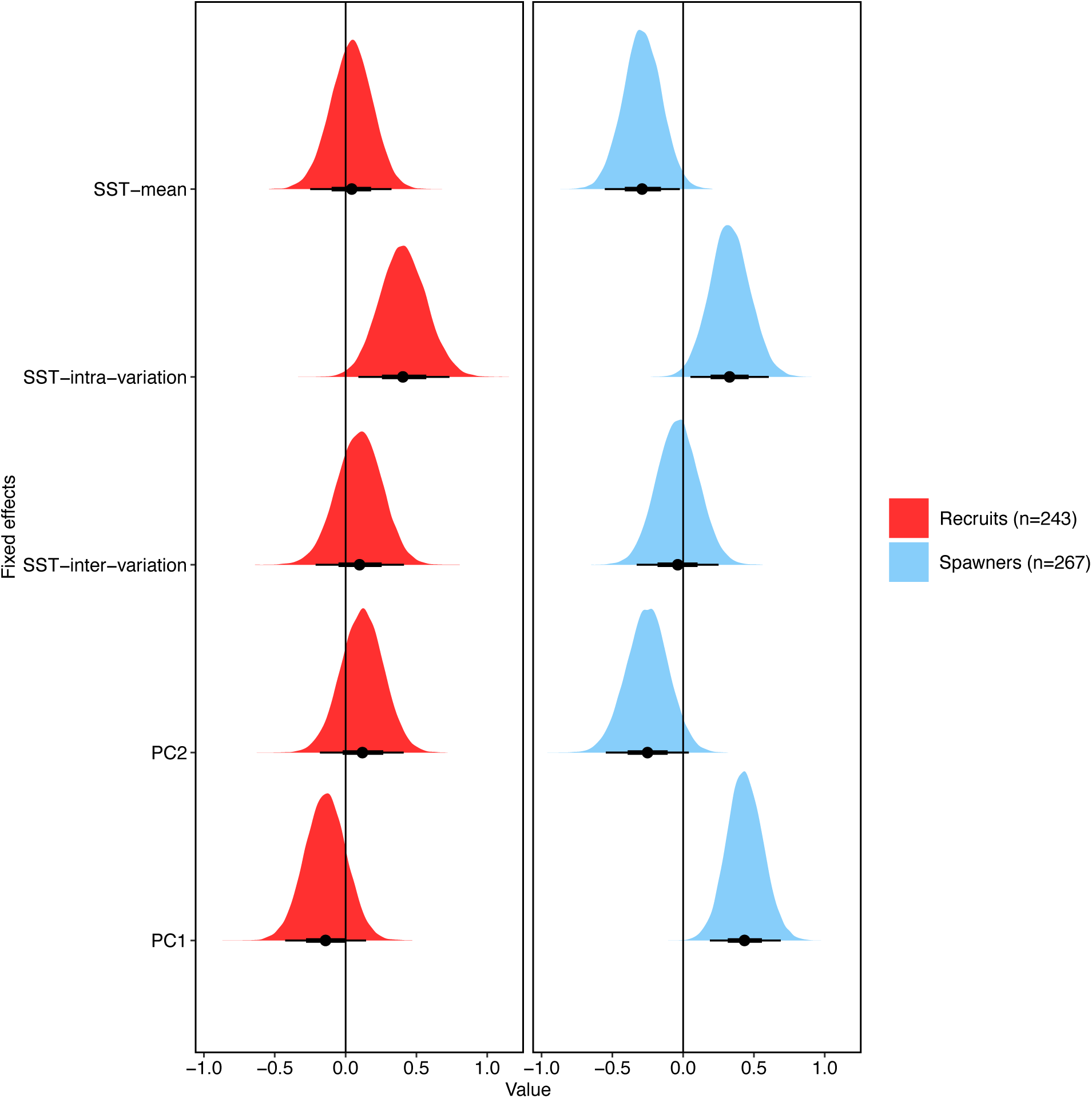
Sensitivity analysis 3 using the value of θ for all stocks that produced the second greatest value of ρ. We present the posterior distributions along with a point estimate for the posterior median and the 66% (thick black line) and 95% (thin black line) credible intervals.

**Fig. S6.**
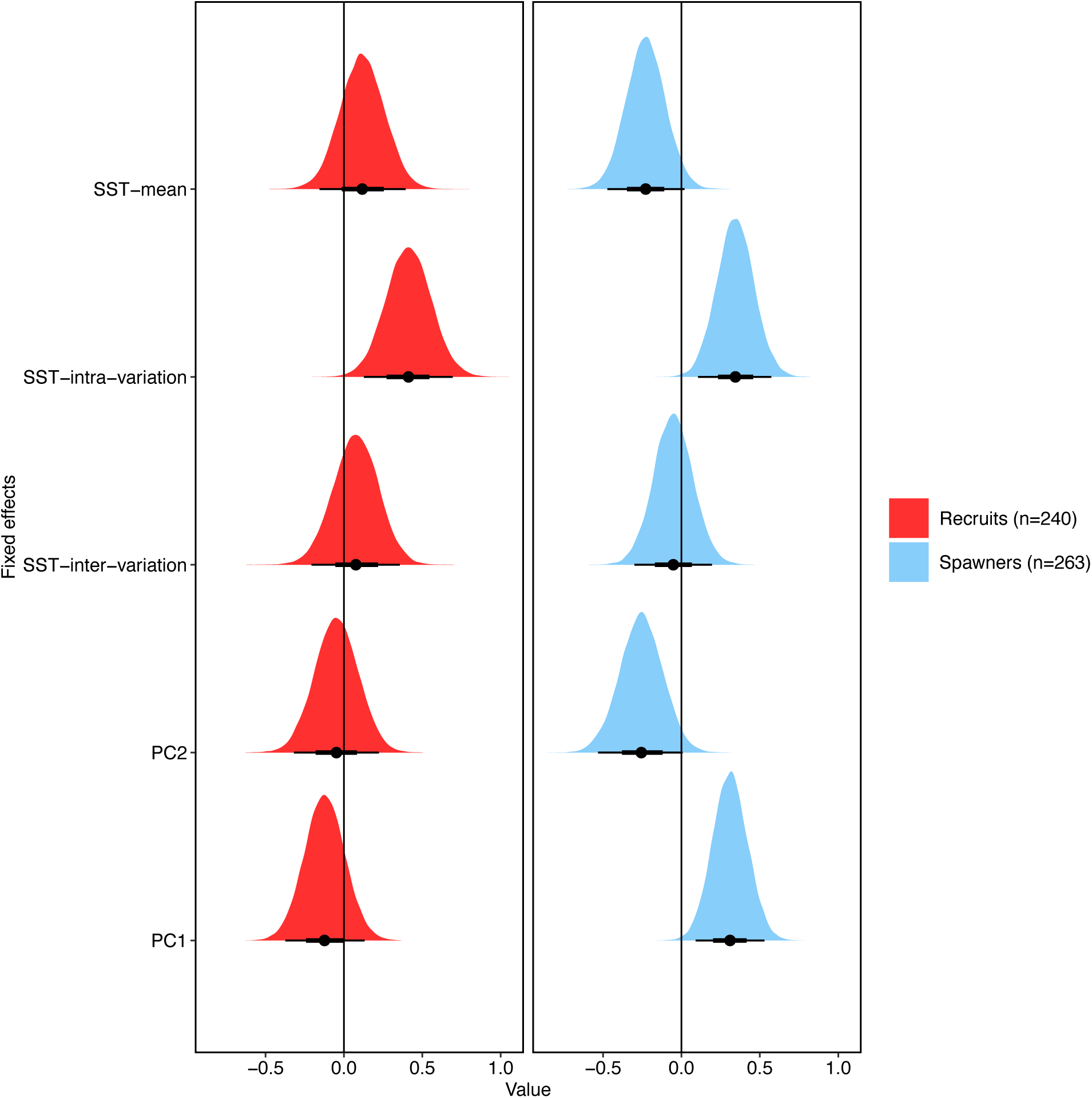
Sensitivity analysis 4 using temperature correlates calculated from the COBE-SST 2 database We present the posterior distributions along with a point estimate for the posterior median and the 66% (thick black line) and 95% (thin black line) credible intervals.

**Fig. S7.**
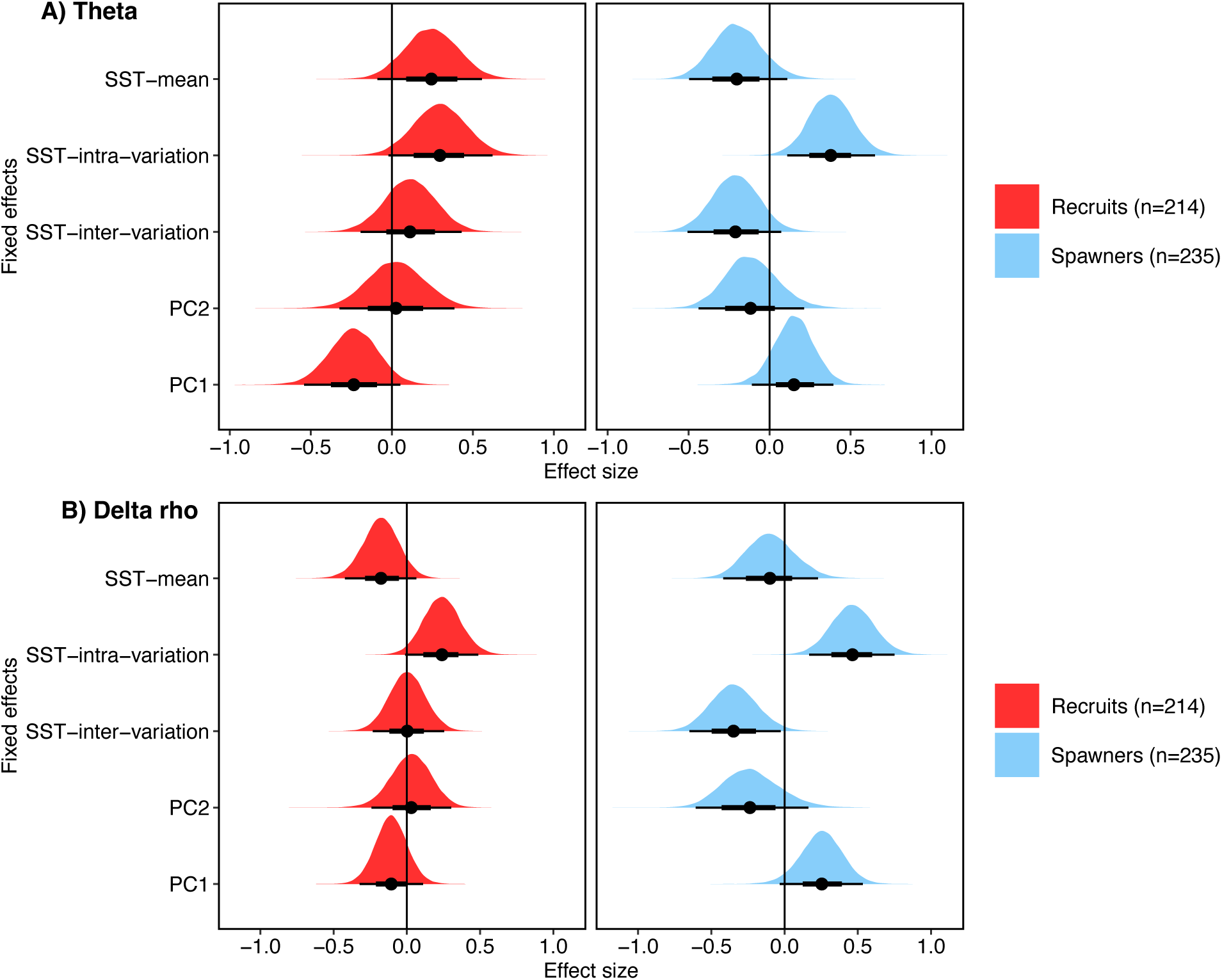
Sensitivity analysis 5 showing that the main results held after correcting for phylogenetic non-independence using a phylogenetic correlation matrix random effect. We present the posterior distributions along with a point estimate for the posterior median and the 66% (thick black line) and 95% (thin black line) credible intervals.

**Fig. S8.**
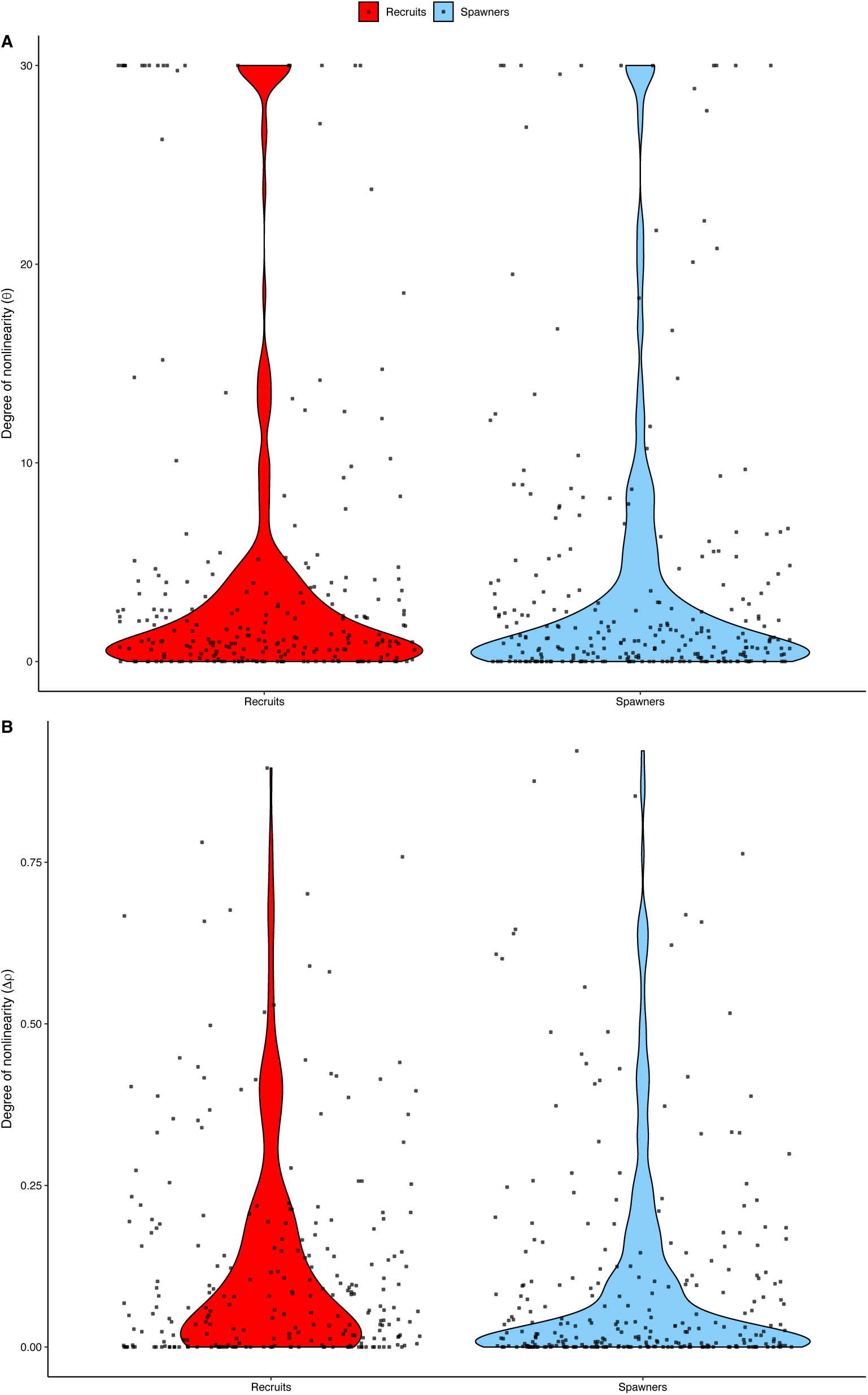
Violin jitter plots showing the distribution of the nonlinearity metrics, θ and Δρ.

**Fig. S9.**
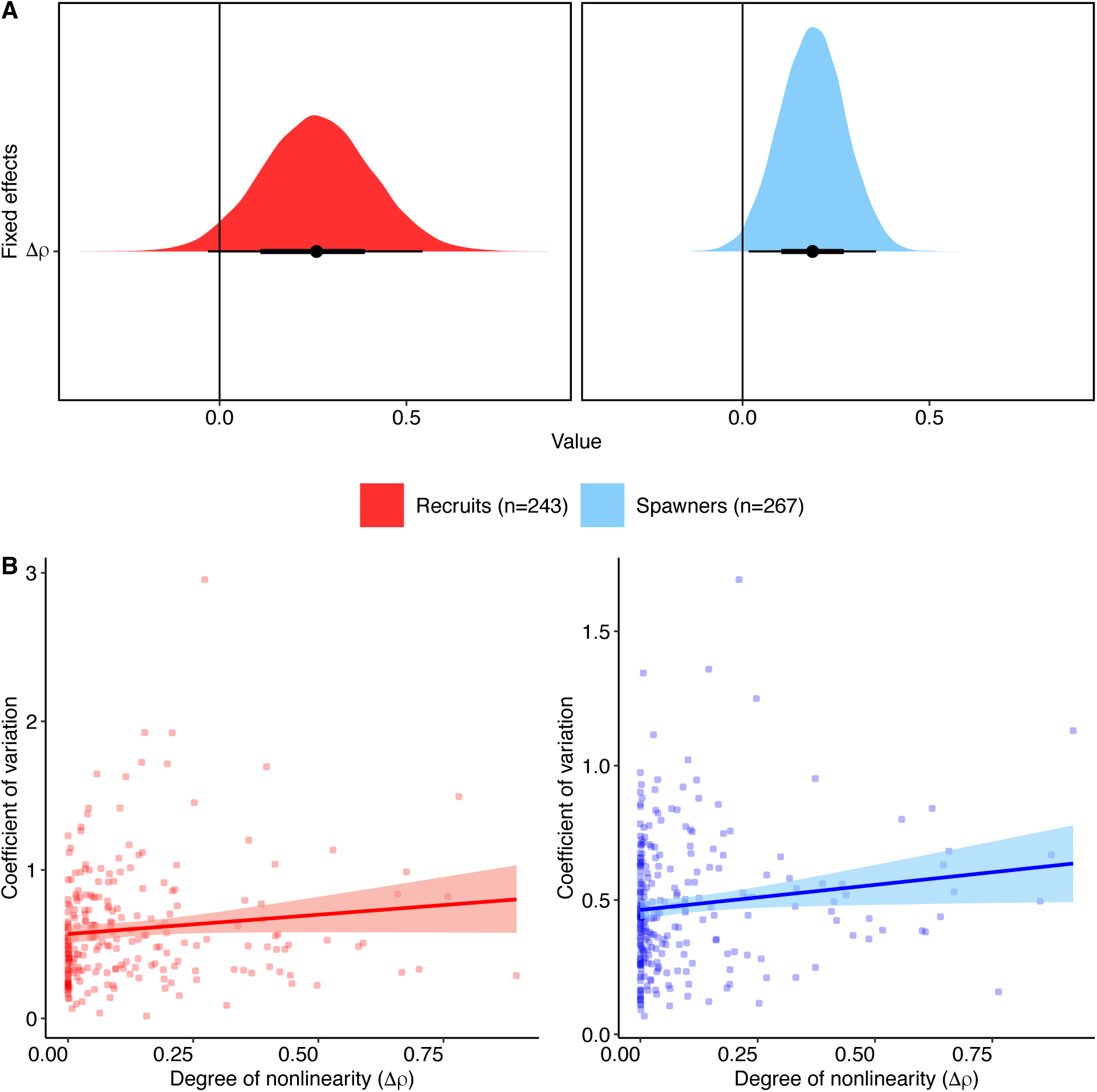
Correlation between nonlinearity (Δρ) and the coefficient of variation (CV) in time series of recruitment and spawner biomass. **A)** Bayesian gaussian models with CV as a function of nonlinearity. We present the posterior distributions along with a point estimate for the posterior median and the 66% (thick black line) and 95% (thin black line) credible intervals. **B)** Conditional effect plots with the line and shaded areas represent the median and 95% credible interval. Red and blue dots represent the values of Δρ for 243 recruit and 267 spawner populations, respectively.

**Fig. S10.**
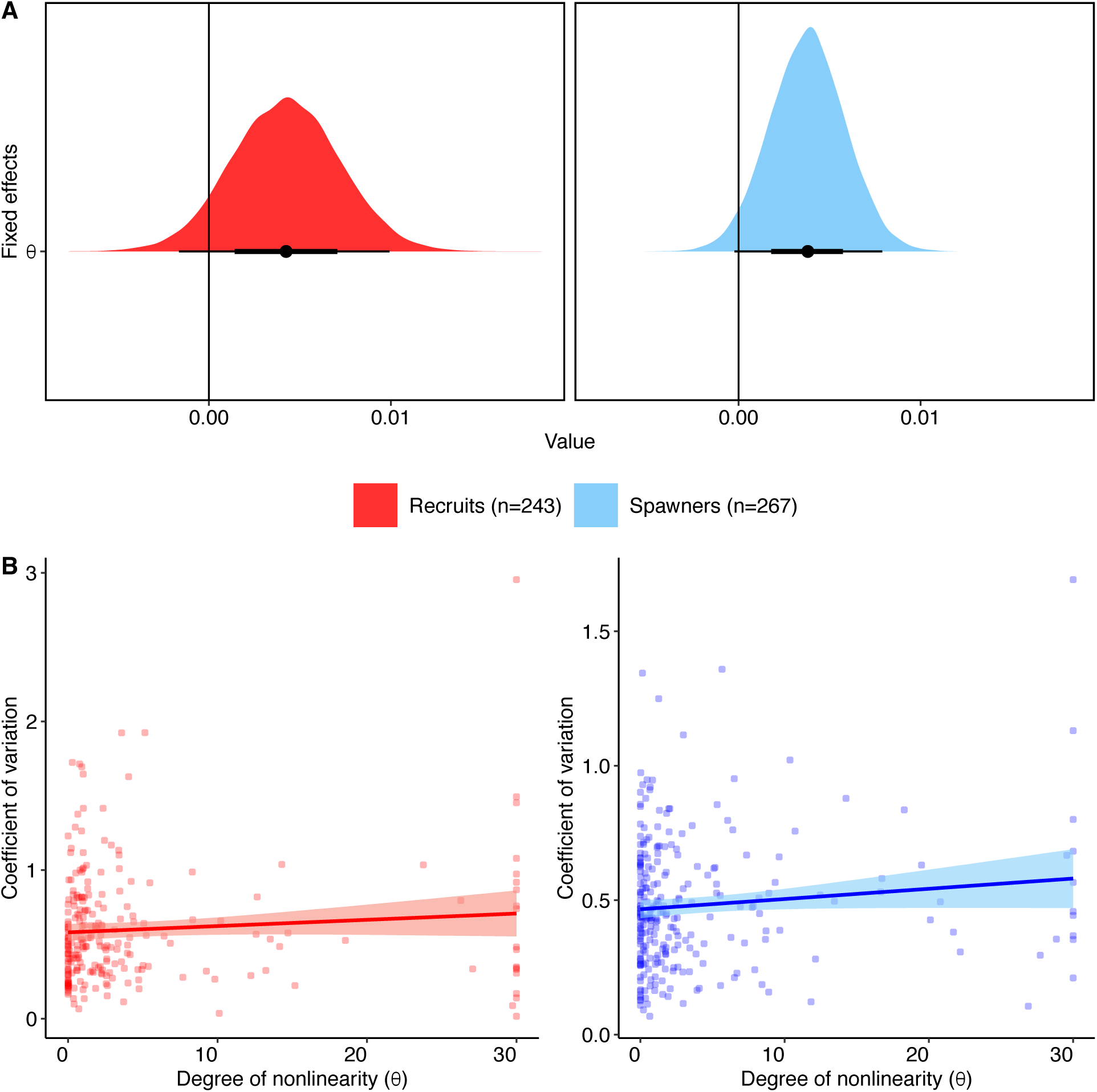
Correlation between nonlinearity (θ) and the coefficient of variation (CV) in time series of recruitment and spawner biomass. **A)** Bayesian gaussian models with CV as a function of nonlinearity. We present the posterior distributions along with a point estimate for the posterior median and the 66% (thick black line) and 95% (thin black line) credible intervals. **B)** Conditional effect plots with the line and shaded areas represent the median and 95% credible interval. Red and blue dots represent the values of θ for 243 recruit and 267 spawner populations, respectively.

**Fig. S11.**
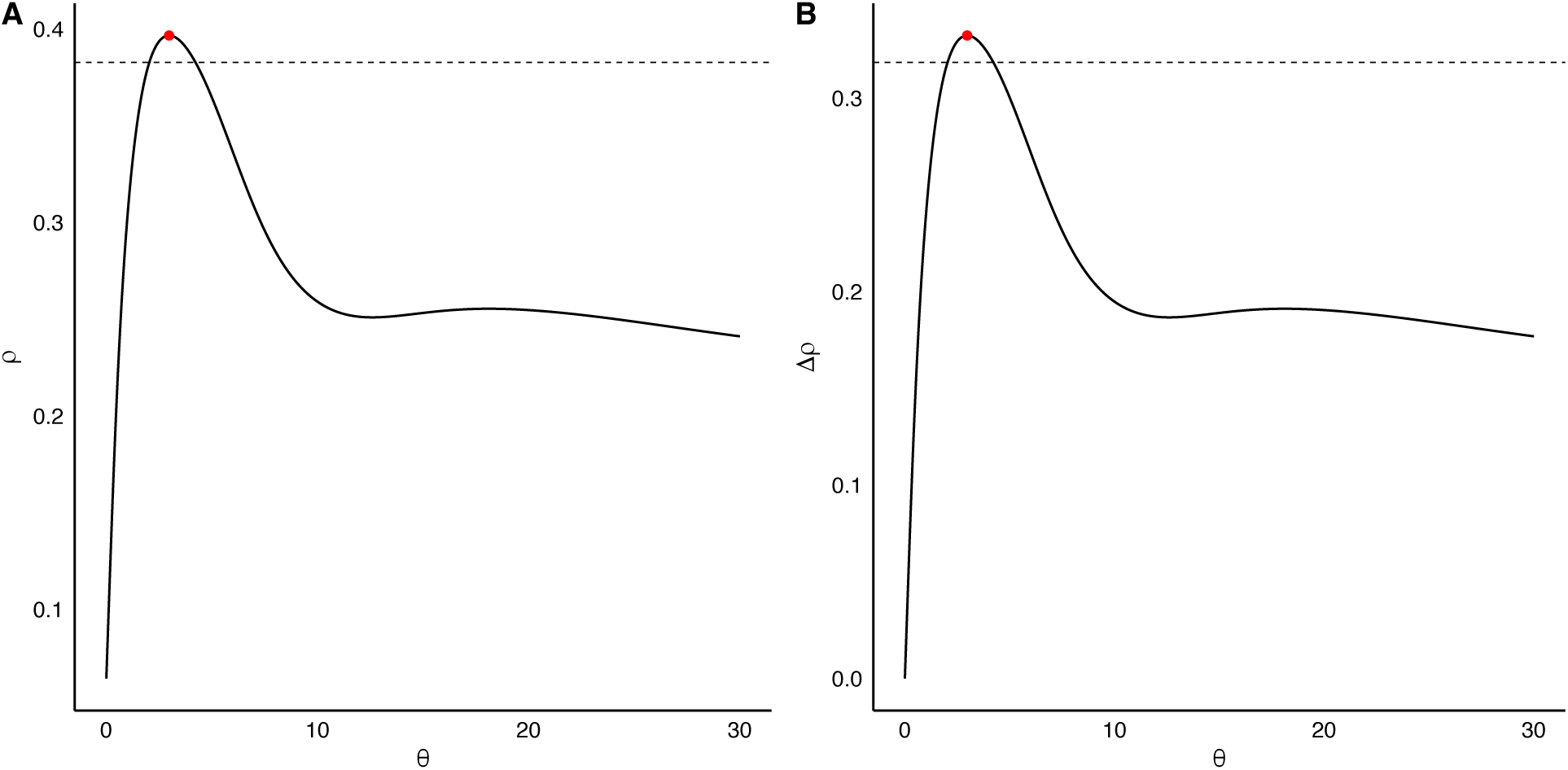
Example case study (spawner population id: RHERRTSST) to demonstrate the threshold for **A)** sensitivity analysis, and **B)** greater uncertainty in θ than Δρ. The red points represent the maximum rho. The horizontal dashed lines represent the 3.5% threshold, and all values of theta corresponding to rho values above this line are sampled from for sensitivity analyses 1 and 2.

## References

1. May, R. M. Simple mathematical models with very complicated dynamics. Nature 261, 459– 467 (1976).

2. Sugihara, G. et al. Are exploited fish populations stable? Proceedings of the National Academy of Sciences 108, E1224–E1225 (2011).

3. Rogers, T. L., Johnson, B. J. & Munch, S. B. Chaos is not rare in natural ecosystems. Nature Ecology & Evolution 6, 1105–1111 (2022).

4. Benincà, E. et al. Chaos in a long-term experiment with a plankton community. Nature 451, 822–825 (2008).

5. Scheffer, M. et al. Early-warning signals for critical transitions. Nature 461, 53–59 (2009).

6. Clark, T. J. & Luis, A. D. Nonlinear population dynamics are ubiquitous in animals. Nature Ecology & Evolution 4, 75–81 (2020).

7. Dakos, V., Glaser, S. M., Hsieh, C. & Sugihara, G. Elevated nonlinearity as an indicator of shifts in the dynamics of populations under stress. Journal of The Royal Society Interface 14, 20160845 (2017).

8. Anderson, C. N. K. et al. Why fishing magnifies fluctuations in fish abundance. Nature 452, 835–839 (2008).

9. Hsieh, C., Glaser, S. M., Lucas, A. J. & Sugihara, G. Distinguishing random environmental fluctuations from ecological catastrophes for the North Pacific Ocean. Nature 435, 336–340 (2005).

10. Berkeley, S. A., Hixon, M. A., Larson, R. J. & Love, M. S. Fisheries Sustainability via Protection of Age Structure and Spatial Distribution of Fish Populations. Fisheries 29, 23–32 (2004).

11. Fischer, E. M. & Knutti, R. Anthropogenic contribution to global occurrence of heavy-precipitation and high-temperature extremes. Nature Climate Change 5, 560–564 (2015).

12. Costantino, R. F., Desharnais, R. A., Cushing, J. M. & Dennis, B. Chaotic Dynamics in an Insect Population. Science 275, 389–391 (1997).

13. Werner, J., Pietsch, T., Hilker, F. M. & Arndt, H. Intrinsic nonlinear dynamics drive single-species systems. Proceedings of the National Academy of Sciences 119, e2209601119 (2022).

14. Beddington, J. R. & May, R. M. Harvesting Natural Populations in a Randomly Fluctuating Environment. Science 197, 463–465 (1977).

15. Hsieh, C. et al. Fishing elevates variability in the abundance of exploited species. Nature 443, 859–862 (2006).

16. Dixon, P. A., Milicich, M. J. & Sugihara, G. Episodic Fluctuations in Larval Supply. Science 283, 1528–1530 (1999).

17. Glaser, S. M. et al. Detecting and forecasting complex nonlinear dynamics in spatially structured catch-per-unit-effort time series for North Pacific albacore (Thunnus alalunga). Canadian Journal of Fisheries and Aquatic Sciences 68, 400–412 (2011).

18. Glaser, S. M. et al. Complex dynamics may limit prediction in marine fisheries. Fish and Fisheries 15, 616–633 (2014).

19. Glaser, S. M., Ye, H. & Sugihara, G. A nonlinear, low data requirement model for producing spatially explicit fishery forecasts. Fisheries Oceanography 23, 45–53 (2014).

20. Munch, S. B., Giron-Nava, A. & Sugihara, G. Nonlinear dynamics and noise in fisheries recruitment: A global meta-analysis. Fish and Fisheries 19, 964–973 (2018).

21. Ye, H. et al. Equation-free mechanistic ecosystem forecasting using empirical dynamic modeling. Proceedings of the National Academy of Sciences 112, E1569–E1576 (2015).

22. Perretti, C. T., Munch, S. B. & Sugihara, G. Model-free forecasting outperforms the correct mechanistic model for simulated and experimental data. Proceedings of the National Academy of Sciences 110, 5253–5257 (2013).

23. Greenman, J. V. & Benton, T. G. The Amplification of Environmental Noise in Population Models: Causes and Consequences. The American Naturalist 161, 225–239 (2003).

24. Shelton, A. O. & Mangel, M. Fluctuations of fish populations and the magnifying effects of fishing. Proceedings of the National Academy of Sciences 108, 7075–7080 (2011).

25. Rogers, L. A., Moore, Z., Daigle, A., Luijckx, P. & Krkošek, M. Experimental evidence of size-selective harvest and environmental stochasticity effects on population demography, fluctuations and non-linearity. Ecology Letters 26, 586–596 (2023).

26. Ricard, D., Minto, C., Jensen, O. P. & Baum, J. K. Examining the knowledge base and status of commercially exploited marine species with the RAM Legacy Stock Assessment Database. Fish and Fisheries 13, 380–398 (2012).

27. Thorson, J. T. et al. Identifying direct and indirect associations among traits by merging phylogenetic comparative methods and structural equation models. Methods in Ecology and Evolution 14, 1259–1275 (2023).

28. Huang, B. et al. Improvements of the Daily Optimum Interpolation Sea Surface Temperature (DOISST) Version 2.1. (2021) doi:10.1175/JCLI-D-20-0166.1.

29. Free, C. M. et al. Impacts of historical warming on marine fisheries production. Science 363, 979–983 (2019).

30. Munch, S. B., Rogers, T. L. & Sugihara, G. Recent developments in empirical dynamic modelling. Methods in Ecology and Evolution 14, 732–745 (2023).

31. Sugihara, G. & May, R. M. Nonlinear forecasting as a way of distinguishing chaos from measurement error in time series. Nature 344, 734–741 (1990).

32. Sugihara, G. Nonlinear Forecasting for the Classification of Natural Time Series. Philosophical Transactions: Physical Sciences and Engineering 348, 477–495 (1994).

33. Sugihara, G. et al. Detecting Causality in Complex Ecosystems. Science 338, 496–500 (2012).

34. Drinkwater, K. F. & Kristiansen, T. A synthesis of the ecosystem responses to the late 20th century cold period in the northern North Atlantic. ICES Journal of Marine Science 75, 2325–2341 (2018).

35. Frank, K. T., Petrie, B., Leggett, W. C. & Boyce, D. G. Towards a more balanced assessment of the dynamics of North Atlantic ecosystems—a comment on Drinkwater and Kristiansen (2018). ICES Journal of Marine Science 76, 2489–2494 (2019).

36. McCann, K., Hastings, A. & Huxel, G. R. Weak trophic interactions and the balance of nature. Nature 395, 794–798 (1998).

37. Frank, K. T., Petrie, B., Fisher, J. A. D. & Leggett, W. C. Transient dynamics of an altered large marine ecosystem. Nature 477, 86–89 (2011).

38. Morris, W. F. et al. Longevity Can Buffer Plant and Animal Populations Against Changing Climatic Variability. Ecology 89, 19–25 (2008).

39. Schindler, D. E. et al. Population diversity and the portfolio effect in an exploited species. Nature 465, 609–612 (2010).

40. Brown, J. H., Gillooly, J. F., Allen, A. P., Savage, V. M. & West, G. B. Toward a Metabolic Theory of Ecology. Ecology 85, 1771–1789 (2004).

41. Roy, K., Jablonski, D., Valentine, J. W. & Rosenberg, G. Marine latitudinal diversity gradients: Tests of causal hypotheses. Proceedings of the National Academy of Sciences 95, 3699–3702 (1998).

42. Hsieh, C., Anderson, C. & Sugihara, G. Extending Nonlinear Analysis to Short Ecological Time Series. The American Naturalist 171, 71–80 (2008).

43. Liu, J. et al. Complexity of Coupled Human and Natural Systems. Science 317, 1513–1516 (2007).

44. Fogarty, M. J., Gamble, R. & Perretti, C. T. Dynamic Complexity in Exploited Marine Ecosystems. Front. Ecol. Evol. 4, (2016).

45. Volkoff, H. & Rønnestad, I. Effects of temperature on feeding and digestive processes in fish. Temperature: Multidisciplinary Biomedical Journal 7, 307 (2020).

46. Himes-Cornell, A. & Kasperski, S. Assessing climate change vulnerability in Alaska’s fishing communities. Fisheries Research 162, 1–11 (2015).

47. Munday, P. L., Jones, G. P., Pratchett, M. S. & Williams, A. J. Climate change and the future for coral reef fishes. Fish and Fisheries 9, 261–285 (2008).

48. Cinner, J. E. et al. Vulnerability of coastal communities to key impacts of climate change on coral reef fisheries. Global Environmental Change 22, 12–20 (2012).

49. Walters, C. J. & Martell, S. J. D. Fisheries Ecology and Management. (Princeton University Press, 2004).

50. Frank, K. T., Petrie, B., Leggett, W. C. & Boyce, D. G. Large scale, synchronous variability of marine fish populations driven by commercial exploitation. Proceedings of the National Academy of Sciences 113, 8248–8253 (2016).

51. Schweigert, J. F., Boldt, J. L., Flostrand, L. & Cleary, J. S. A review of factors limiting recovery of Pacific herring stocks in Canada. ICES Journal of Marine Science 67, 1903– 1913 (2010).

52. Giron-Nava, A. et al. Environmental variability and fishing effects on the Pacific sardine fisheries in the Gulf of California. Can. J. Fish. Aquat. Sci. 78, 623–630 (2021).

53. Tsai, C.-H., Munch, S. B., Masi, M. D. & Stevens, M. H. Empirical dynamic modeling for sustainable benchmarks of short-lived species. ICES Journal of Marine Science 81, 1209– 1220 (2024).

54. Walters, C. J. & Juanes, F. Recruitment Limitation as a Consequence of Natural Selection for Use of Restricted Feeding Habitats and Predation Risk Taking by Juvenile Fishes. Canadian Journal of Fisheries and Aquatic Sciences 50, 2058–2070 (1993).

55. Andersen, K. H., Jacobsen, N. S., Jansen, T. & Beyer, J. E. When in life does density dependence occur in fish populations? Fish and Fisheries 18, 656–667 (2017).

56. Myers, R. A. & Cadigan, N. G. Density-Dependent Juvenile Mortality in Marine Demersal Fish. Canadian Journal of Fisheries and Aquatic Sciences 50, 1576–1590 (1993).

57. Hechler, R. & Krkosek, M. Data and code for: Temperature variation and life history drive nonlinear dynamics of marine fish populations. OSF (2025).

58. COBE-SST 2 and Sea Ice: NOAA Physical Sciences Laboratory COBE-SST 2 and Sea Ice. https://psl.noaa.gov/data/gridded/data.cobe2.html.

59. Thorson, J. T. Predicting recruitment density dependence and intrinsic growth rate for all fishes worldwide using a data-integrated life-history model. Fish and Fisheries 21, 237–251 (2020).

60. SugiharaLab/rEDM: Applications of Empirical Dynamic Modeling from Time Series. https://github.com/SugiharaLab/rEDM.

61. Ye, H., Deyle, E. R., Gilarranz, L. J. & Sugihara, G. Distinguishing time-delayed causal interactions using convergent cross mapping. Sci Rep 5, 14750 (2015).

62. Bürkner, P.-C. brms: An R Package for Bayesian Multilevel Models Using Stan. Journal of Statistical Software 80, 1–28 (2017).

63. R: The R Project for Statistical Computing. https://www.r-project.org/.

64. Morey, R. BayesFactor: Computation of Bayes Factors for Common Designs. (2023).

65. Gelman, A. & Rubin, D. B. Inference from Iterative Simulation Using Multiple Sequences. Statistical Science 7, 457–472 (1992).

66. Rabosky, D. L. et al. An inverse latitudinal gradient in speciation rate for marine fishes. Nature 559, 392–395 (2018).

